# Private information leakage from functional genomics data: Quantification with calibration experiments and reduction via data sanitization protocols

**DOI:** 10.1101/345074

**Authors:** Gamze Gürsoy, Prashant Emani, Charlotte M. Brannon, Otto A. Jolanki, Arif Harmanci, J. Seth Strattan, Andrew D. Miranker, Mark Gerstein

**Affiliations:** Program in Computational Biology and Bioinformatics, Yale University, New Haven, CT 06520, USA; Department of Molecular Biophysics and Biochemistry, Yale University, New Haven, CT 06520, USA; Stanford University School of Medicine, Department of Genetics, Stanford, CA 94305, USA; School of Biomedical Informatics, Center for Precision Health, University of Texas Health Sciences Center, Houston TX, 77030, USA; Department of Chemical and Environmental Engineering, Yale University, New Haven, CT 06520, USA; Department of Computer Science, Yale University, New Haven, CT 06520, USA

**Keywords:** genome privacy, functional genomics, linkage attacks, surreptitious DNA sequencing, data sanitization

## Abstract

The generation of functional genomics datasets is surging, as they provide insight into gene regulation and organismal phenotypes (e.g., genes upregulated in cancer). The intention of functional genomics experiments is not necessarily to study genetic variants, yet they pose privacy concerns due to their use of next-generation sequencing. Moreover, there is a great incentive to share raw reads for better analyses and general research reproducibility. Thus, we need new modes of sharing beyond traditional controlled-access models. Here, we develop a data-sanitization procedure allowing raw functional genomics reads to be shared while minimizing privacy leakage, thus enabling principled privacy-utility trade-offs. It works with traditional Illumina-based assays and newer technologies such as 10x single-cell RNA-sequencing. The procedure depends on quantifying the privacy leakage in reads by statistically linking study participants to known individuals. We carried out these linkages using data from highly accurate reference genomes and more realistic environmental samples.

## 1 Introduction

Advances in sequencing technologies and laboratory techniques have enabled researchers to comprehensively probe epigenetic and transcriptomic states of the cell, such as gene expression levels or DNA-binding protein levels, the majority of which are clinically actionable [e.g., The Cancer Genome Atlas (TCGA)]. With the availability of more advanced techniques, such as single-cell RNA sequencing (scRNA-Seq) and single-cell assay for transposase-accessible chromatin using sequencing, we can now functionally annotate tissues even at the single-cell level. This increased resolution will soon bring a surge of new functional genomics datasets from large cohorts of individuals, surpassing the available sequenced genomes. For example, let us consider a human biosample being assayed for all aspects of its biology. We can sequence the DNA of the sample once, but numerous functional genomics assays can be performed on the same sample. Moreover, as more data are collected from a larger number of studied individuals, we can boost statistical power for future discoveries–that is, if the collected data are made available to the wider research community.

Privacy is a barrier to the open sharing of functional genomics datasets, as individuals’ genetic variants can be inferred from the data. Studies on genomic privacy have traditionally focused on DNA, partly due to the dominance of DNA-sequencing data in genomics research [1, 2, 3, 4, 5, 6]. However, as the balance in data acquisition shifts toward large-scale functional genomics data, privacy studies must shift as well [7]. From a privacy standpoint, functional genomics data have a two-sided nature, in contrast to DNA-sequencing data. Although these experiments are usually performed to understand the biology of cells, rather than to reveal identifying genetic variants, raw reads are nonetheless tagged by the genetic variants of the individuals due to the experimental design. Moreover, in addition to including genetic variant information that can be inferred from raw reads, functional genomics data provide the conditions under which the assay was performed (e.g., linking a specific phenotype to the sample). Furthermore, lower barriers to sequencing data enable a wider range of people, including “citizen scientists,” to access genomic data that can be overlapped with functional genomics data to infer sensitive information about study participants.

These privacy concerns should be addressed while considering the unique aspects of functional genomics data in order to maximize open sharing. To answer critical biological questions, researchers must generate key properties from functional genomics data, such as gene expression quantifications or transcription factor (TFF) binding enrichments. These properties must be generated from the raw reads; however, such calculations do not require genetic variants in the reads. Sharing the raw reads, as opposed to derived quantities, is essential because this sharing provides researchers with the necessary level of control over their analyses and helps advance biomedical science by permitting a rapid assessment of tools and methods, thus enhancing reproducibility. Currently, large consortia such as TCGA [8], the Genotype-Tissue Expression (GTEx) project [9], and PsychENCODE [10] store raw functional genomics reads behind controlled access following traditional DNA-sharing protocols, while providing open access to summary-level data such as gene expression levels. This approach makes sense; however, these large consortia and other researchers are aware that a number of genotypes can be inferred from summary-level data [11, 12, 13]. Thus, the scale of the genotype leakage from functional genomics reads is much larger, far beyond an acceptable level. Meanwhile, controlled-access sharing protocols delay data access for average researchers by creating bureaucratic bottlenecks or technical challenges. Moreover, these protocols were developed with a focus on DNA-sequencing data, which are obtained for the purpose of identifying the genetic variants of an individual. Thus, a new model for sharing raw functional genomics reads is needed, one that better fits the privacy and utility needs of functional genomics data.

In this study, we developed read-sanitization techniques that enable public sharing of minimally manipulated functional genomics reads, while protecting sensitive information and minimizing the amount of private data requiring special access and storage. Our methods are based on an optimized balance between privacy requirements and the inherent utility of functional genomics reads. Our aim is to provide reads to the community that can be easily used as inputs to any functional genomics data processing pipeline.

To develop data-sanitization techniques to reduce and even eliminate privacy risk, a comprehensive assessment of private information leakage from different types of functional genomics reads under varying noise conditions and sequencing coverage levels is necessary. Accordingly, we performed common privacy breaches using publicly available functional genomics and genome sequencing datasets. To quantify the privacy loss in a more realistic setting, we conducted DNA-sequencing and RNA-sequencing (RNA-Seq) assays on blood tissue and environmental samples, such as used coffee cups (to mimic surreptitious DNA testing) collected from consented individuals. We demonstrated that our statistical measures can link individuals to functional genomics datasets under different noise levels and sequencing coverage levels, hence revealing sensitive phenotype information about the individuals. We then showed that our data-sanitization protocol minimizes this privacy risk, while providing accurate estimations of key summary quantities, such as gene expression levels. Finally, we demonstrated that easy access to a large amount of data that are otherwise locked behind controlled access can be achieved with a principled balance between privacy and utility.

## 2 Results

### 2.1 Privacy Breach Scenarios for Quantification of Privacy Loss

A “linkage attack” is a common breach of privacy in which one can quantify the private information leakage in an anonymized dataset 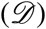 by using publicly available information (*ℐ*) about the individuals in the dataset [14, 1]. A famous example of a linkage attack is the de-anonymization of Netflix Prize challenge data 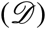 using publicly available IMDB user comments (*ℐ*) [14]. Within genomics, researchers have shown that genetic information from publicly available genealogy websites (*ℐ*) can be overlapped with short tandem repeats on the Y chromosomes of anonymized genomes to infer the surnames of participants in genomics datasets 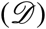 [6]; likewise, expression quantitative trait locus (eQTL) data (*ℐ*) can be applied to de-anonymize gene expression datasets 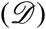 [11, 12, 13]. Linkage techniques have been used outside of the privacy context as well, such as to resolve sample swap problems during omics data production [15, 16, 17, 18].

Here, we define three types of linkage attacks (Figure S1), which differ by the nature of the data in 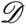 and *ℐ*, i.e., whether the data is perfect (P) or noisy (N):

- (I) Case P-P involves obtaining publicly available perfect information *ℐ* about a known individual (e.g., date of birth, zip code, gender) and overlapping it with perfect information in anonymized dataset 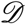 (e.g., zip code, gender) to reveal sensitive information (e.g., political affiliation). This can be done by cross-referencing two datasets, and often does not require any statistical heuristics. A famous example of this kind of attack in medicine is the use of cross-referencing birthdate and zip code information present in medical records and voter list datasets to reveal the addresses of patients [19].
- (II) Case P-N involves obtaining publicly available perfect information *ℐ* about a known individual (e.g., whole-genome sequencing reads) and overlapping it with noisy information from anonymized dataset 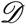 (e.g., ChIP-Seq reads) to reveal sensitive information (e.g., psychiatric disease status). Because this type of attack involves linking information to a noisy dataset, simple overlaps often do not work and statistical analysis is required.
- (III) Case N-N involves obtaining publicly available noisy information *ℐ* about a known individual [e.g., whole-genome sequencing (WGS) of DNA gathered from a coffee cup] and overlapping it with noisy information from anonymized dataset 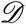 (e.g., ChIP-Seq reads) to reveal sensitive information (e.g., psychiatric disease status). This can require even more involved statistical techniques to sort through the signal in two noisy datasets.

Thus, linkage becomes more difficult with noisier data as it requires statistical techniques rather than simple overlaps. In order to quantify the private information leakage in functional genomics datasets, we adopted Narayanan and Shmatikov’s statistical techniques of privacy breaches [14]. The privacy threat models are as follows. One can imagine a scenario in which an adversary gains illicit access to a known individual’s WGS data and performs linkage attacks on functional genomics data in order to infer private phenotypes associated with them. Note that, in this scenario, the private information that will be leaked are sensitive *phenotypes* (e.g., HIV status, bipolar disorder status, etc.), not *genotypes*. The genotypes in the functional genomics reads are merely used as a means to infer the phenotype. This is somewhat different from traditional privacy attacks, in which the leaked information is the genotype data. The genotypes obtained from functional genomics data are noisy, functional genomics reads constitute a noisy dataset 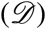, and genomes can be thought of as perfect information (*ℐ*) about the individual. Therefore, we mimicked this scenario with a Case P-N attack. One can also imagine a more realistic, “real-world” scenario: an adversary takes advantage of the DNA trail individuals leave behind in everyday life, such as saliva on a stamp [20], a used facial tissue, or a morning cup of coffee. As in the previous example, the genotypes obtained in this case are noisy, but the DNA information (*ℐ*) extracted from real-world samples (e.g., coffee cups) is also noisy due to possible exposure to multiple individuals other than the owner. Therefore, we mimicked this scenario with a Case N-N attack. To do so, we performed DNA sequencing and RNA-Seq assays on blood tissue and coffee-cup samples collected from consented individuals.

### 2.2 Sensitive phenotypes can reliably be inferred by linking perfect genotypes to noisy genotypes called from functional genomics data

Let us assume a study that aims to understand the changes in gene expression of individuals with bipolar disorder (BPD). This study assays the gene expression of a cohort of individuals with BPD-positive (BPD+) and −negative (BPD−) phenotypes and publicly releases the RNA-Seq reads (dataset 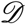). By lawful or unlawful means, an adversary obtains access to the WGS or genotyping array data (information *ℐ*) of a known individual (henceforth referred to as the query individual) and aims to predict whether this individual’s phenotype is BPD+ or BPD−.

The adversary uses traditional variant callers (e.g., GATK [21, 22]) to obtain genotypes from the raw reads of the functional genomics (RNA-Seq) experiments. They create a database of genotypes *S^D^* using raw reads from all the individuals in the study (dataset 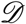). The genotypes obtained from information *ℐ* of the query individual are denoted as 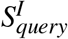. *S^D^* contains partial (i.e., missing some alleles) and noisy (i.e., containing some misidentified alleles) genotypes, as genotyping from RNA-Seq reads may contain errors and cannot cover the entire genome. By the design of the functional genomics study, *S^D^* is linked to the phenotype of the individuals (i.e., BPD+ and BPD−). The adversary then statistically matches the genotypes of the query individual, 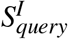, to the genotypes of the individuals, *S^D^*, as follows (see also Figure 1a and Methods): The adversary first calculates a “linking score” as the sum of the log of the inverse of the genotyping frequency of each genotype at the intersection of 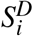 and 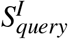 for each entry in *S^D^*. The adversary then ranks the linking scores and calculates a *gap* value for the top-ranked entry in *S^D^* by determining the ratio between the highest and second-highest linking scores. The *gap* value is then compared against *gap* values obtained using a random set of genotypes to assess the statistical significance of the real *gap* value. Finally, the adversary denotes the best match as the query individual and this reveals the BPD status of the known query individual to the attacker. This linking approach (see Methods for mathematical details) is adopted and modified from the Netflix attack by Narayanan and Shmatikov [14].

**Figure 1:**
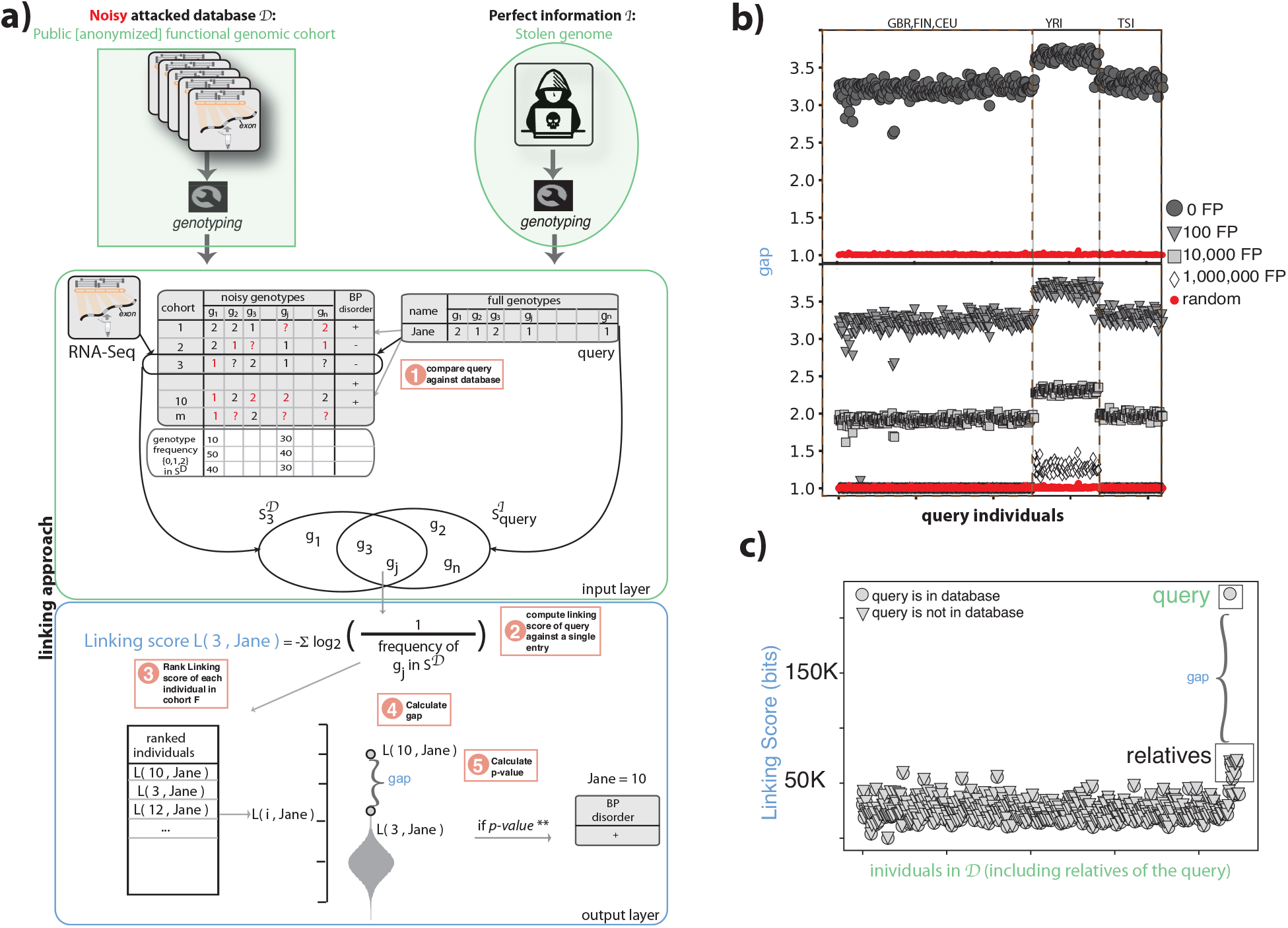
**(a)** Anonymized functional genomics data from a cohort of individuals can be seen as a dataset 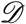 to be attacked. This dataset 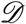 contains functional genomic reads and phenotypes for every individual in the cohort. The perfect information *ℐ* about an individual can be assumed to be WGS data for an individual with known identity. The goal is to determine the phenotype of the known individual by statistically linking the WGS data to the functional genomics cohort. To do so, the attacker performs variant calling on the functional genomics reads and WGS reads. Each individual in the cohort is then scored based on the frequency of the overlapping genotypes between the known individual and the anonymized individual in the cohort. These scores are then ranked and the top-ranked anonymized individual in the cohort is selected as the known individual if the ratio between the top and second-best score (*gap*) is greater than the *gap* values obtained using random sets of genotypes as perfect information *ℐ*. **(b)***gap* values for the 1,000 Genomes individuals in the gEUVADIS RNA-Seq cohort. We included a total of 421 individuals with WGS (from 1,000 Genomes Project) and functional genomics data (from gEUVADIS data). We successfully linked of these individuals to the correct individuals in the cohort with statistical confidence. Red circles are the gap values obtained by linking a random set of genotypes to the phenotype panel. *gap* values are also shown after adding false-positive genotypes to the genotype set of each individual in dataset 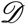. As the total number of false-positive genotypes increases, the *gap* values converge to random. **(c)** The linking scores for each individual in the functional genomics cohort 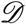 after the addition of genetically related individuals to the queried individual (NA12878). Two scenarios were tested: (i) the functional genomics data of queried individual NA12878 when present in dataset 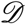 and (ii) the functional genomics data of queried individual NA12878 when not present in dataset 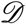. The former showed that addition of genetically related individuals to dataset 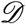 does not affect the linking ability. The latter showed adding genetically related individuals to dataset 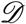 does not cause one of the relatives to be misidentified as the queried individual.

In this study, we used RNA-Seq data of 421 individuals from the gEUVADIS project [23] as the functional genomics dataset 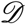, and high-coverage WGS of the same individuals from the 1,000 Genomes Project [24] as the information *ℐ* (Figure 1a). After creating the genotype panel *S^D^* and query genotypes *S^I^*, we successfully linked all 421 query individuals to dataset 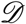 and revealed sensitive phenotypes with a *p*-value < 10^−2^. The top panel of Figure 2b shows the gap values for the best matches. We also generated random sets of genotypes for each query individual and calculated random *gap* values to assess the statistical significance of the true *gap* values (see Methods for details on the statistical significance). This analysis substantiates the reliability of the linking.

**Figure 2:**
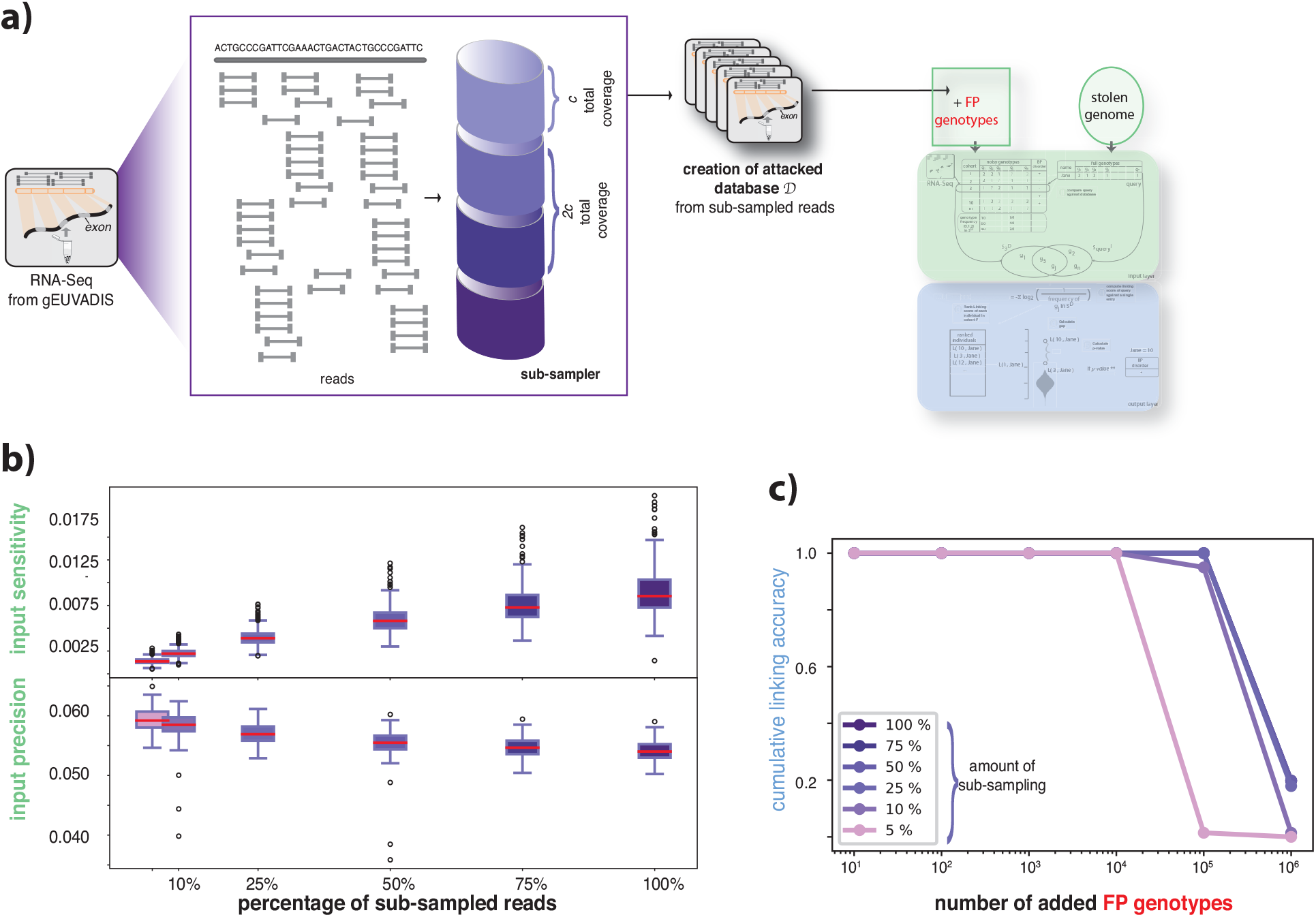
**(a)** The process of pooling reads from functional genomics data to generate different attacked datasets 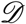 using the subsampled reads. *c* amount of reads were pooled and a new cohort was generated. A linkage attack was performed for each new dataset 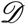 using WGS data as perfect information *ℐ*. **(b)** Variant calling and genotyping were performed for pooled reads for each individual in the dataset 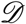. These genotypes were then compared against the gold-standard genotypes obtained from WGS to calculate the precision and sensitivity at each sub-sampling percentage for all individuals. The panel shows the distribution of precision and sensitivity over all individuals in the dataset 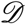 at different subsampling percentages. Precision and sensitivity allow us to understand the noise levels in the functional genomics data. **(c)** Cumulative linking accuracy before and after adding the false-positive genotypes. Linking accuracy was calculated as the ratio between the total number of correctly linked individuals with *p*-values < 0.01 and the total number of queried individuals. This calculation was repeated for each subsampled dataset 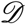.

To further increase the noise levels in the attacked dataset 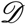, we added an increasing number of randomly picked false-positive genotypes to each genotype call set 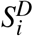 and kept the number of true positive variants constant (see Methods for details). We successfully and significantly linked 418 out of 421 individuals to the cohort even after adding 100,000 false-positive variants to the dataset (Figure 1b). The *gap* values become non-significant only after adding one million false-positive variants (nearly a quarter of the total number of variants in an individual genome, Figure 1b). Interestingly, the RNA-Seq data from individuals with African ancestry (denoted as YRI in Figure 1b), despite having the same coverage as the rest of the RNA-Seq data (from European ancestries, i.e., GBR, FIN, CEU, and TSI), were more vulnerable to genotype-based linkage attacks (Figure 1b). This result is likely due to a higher number of heterozygous or homozygous alternative alleles in the African genomes compared to the reference genome.

#### Effect of genetically related individuals

We next assessed the effect of including related individuals in the attacked dataset 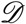. We genotyped the RNA-Seq reads obtained from 14 individuals genetically related to the 1,000 Genomes individual NA12878, including the parents and children [25], and added them to the genotype database *S^D^*. The goal was to determine whether any of the related individuals would be picked as the best match during an attack. We first used the NA12878 genome as the query 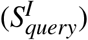 without including NA12878 RNA-Seq in the attacked dataset 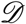. We found that linking scores for the related individuals were very similar to the other individuals in the database; therefore, the *gap* values were not significant and none of the related individuals were mistakenly picked as the query (NA12878). By contrast, when we added the RNA-Seq data of NA12878 to the attacked database, we could significantly match the query genotype set to the correct RNA-Seq data (Figure 1c). Thus, the linking approach used in this study is robust to the addition of even first-degree genetically related individuals to the attacked database. The reason for this specificity is that the linking score depends on the genotypes rather than the existence of the variants. Although first-degree relatives (e.g., mother and son) share a great number of variants, the genotypes of these variants can differ in related individuals.

#### Effect of sequencing depth

The linking schemes described above were done using all the reads in the RNA-Seq dataset. However, the number of reads in a typical functional genomics dataset can vary depending on various factors such as the study design, number of assayed samples, assay type, or tissue type. These variations can greatly affect the number and quality of the genotypes that can be called from the functional genomics data. To understand the effect of the sequencing coverage on the noise levels of the attacked database and on the linking accuracy, we subsampled reads from all the individuals in the gEUVADIS cohort and created cohort genotype datasets *S^D^* for each level of subsampling by using 5, 10, 20,…, up to 100% of the reads in each RNA-Seq experiment (see Figure 2a for a depiction of the subsampling procedure). We first calculated the sensitivity and precision of the called genotypes by using the genotypes from high-coverage WGS data as the ground truth. We calculated the sensitivity as the ratio between the called true positive genotypes and the total (ground-truth) genotypes that belong to the individual. For precision, we calculated the ratio between the called true positive genotypes and all called genotypes. As shown in Figure 2b, the sensitivity (i.e., true positive rate) of the called genotypes improves as we add more reads. However, the sensitivity is still quite low (∼0.75%) even when using all of the reads in an experiment, suggesting that the attacked database missed many of the correct genotypes. By contrast, precision showed a trend towards becoming worse with added reads (from 6% to 5.5%, Figure 2b). Similar to sensitivity, precision was also low even when we used all of the reads in an experiment, suggesting that the attacked database contains several false-positive genotypes and the number of false positives only increases when more reads are sequenced. We then used these databases created from subsampled reads as the attacked databases 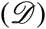 and queried them using the perfect genotypes using the approach outlined above. We performed the linking by a) using the genotypes obtained from reads only under different coverages and b) adding random false-positive variants (ranging from ten to one million) to the genotypes of each entry in the attacked database. We found that linking accuracy, unlike genotyping accuracy, was not affected by the sequencing depth even when we added 10,000 random false-positive genotypes to the entries, as we were able to correctly match all 421 query individuals to the correct RNA-Seq entries in 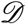.

### 2.3 Sensitive phenotypes can be reliably inferred by linking surreptitiously gathered, noisy genotypes to noisy genotypes called from functional genomics data

Let us assume that the adversary is not able to access existing WGS or genotyping array datasets, and instead obtains environmental DNA samples (e.g., used coffee cups). The adversary aims to identify the disease status of the coffee cups’ owners by linking their DNA to a functional genomics cohort. To mimic this scenario, we performed RNA-Seq experiments on blood tissues from two consented individuals. We then combined these raw RNA-Seq reads with the gEU-VADIS RNA-Seq cohort to create the noisy attacked dataset 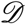. We performed variant calling and obtained sets of genotypes for each individual 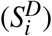. We then collected six used coffee cups from the same individuals. After extracting and amplifying the DNA from the surface of coffee-cup lids using known forensic techniques with commercially available kits (see Methods for details), we performed WGS experiments at 10x sequencing coverage. Each of these WGS experiments was used as the noisy information *ℐ*. We performed variant calling to these low-coverage WGS data from coffee cups (See Tables S1&S2 for mapping and genotyping statistics). Using the linking approach detailed above, we calculated the *gap* values for each query individual by intersecting the genotypes obtained from a coffee cup 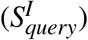 with the genotypes called from RNA-Seq reads for each individual *ℐ* in the attacked dataset 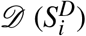 (Figure 3a).

**Figure 3:**
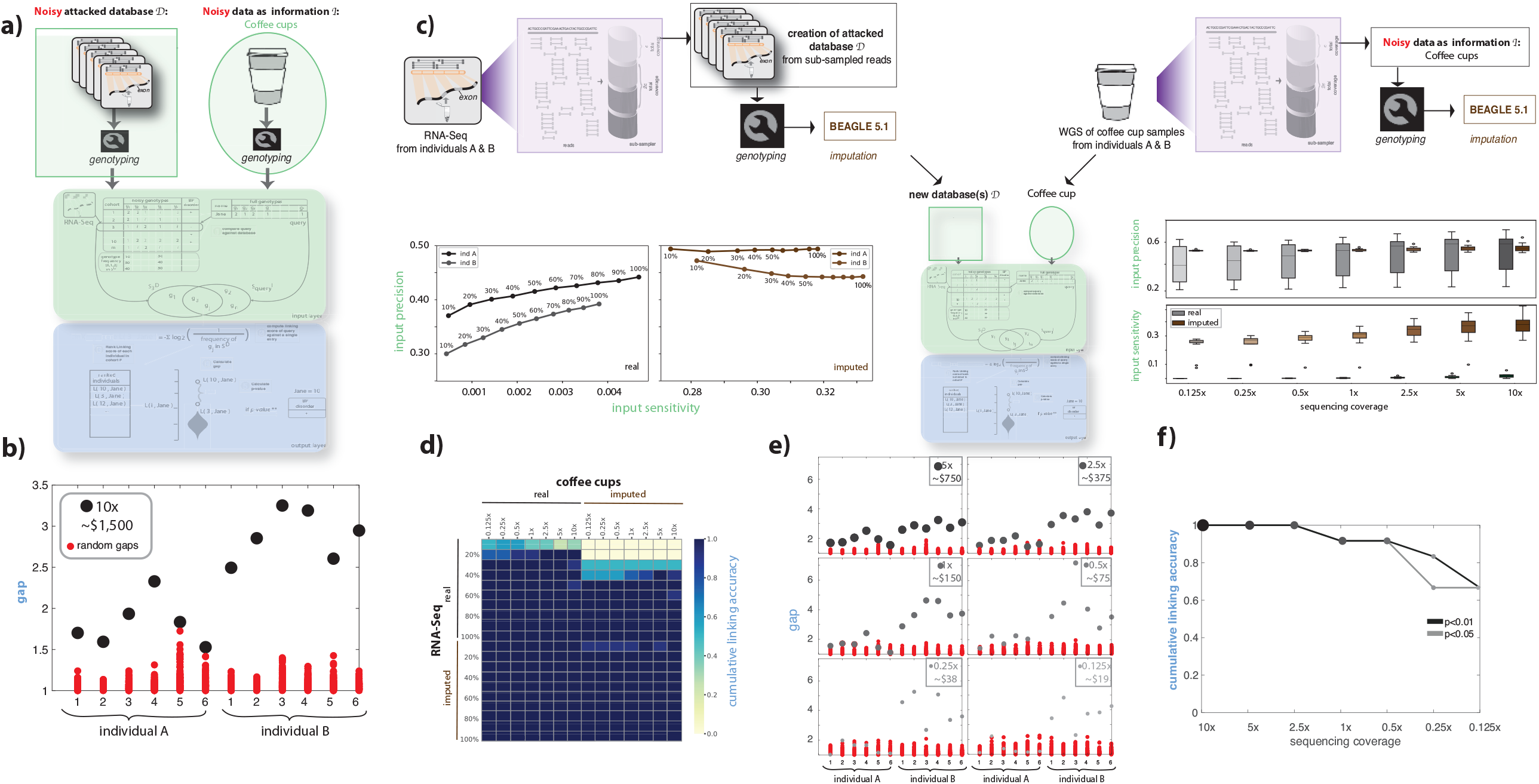
**(a)** Anonymized functional genomics data from a cohort of individuals can be seen as a dataset 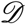 to be attacked. This dataset 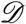 contains functional genomic reads and phenotypes for every individual in the cohort. The noisy information *ℐ* about an individual can be assumed to be WGS data from a surreptitiously obtained environmental sample (e.g., a used coffee cup) of an individual with known identity. The goal is to determine the phenotype of the known individual by statistically linking the coffee cup data to the functional genomics cohort using the statistics described in Figure 1a. **(b)***gap* values for two individuals and six coffee cups, each at 10x sequencing coverage. Each of the coffee cups was successfully linked to its owner in the functional genomics cohort with *p* values< 0.01. **(c)** The process of generating different datasets 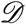 and noisy information *ℐ* by (i) subsampling the reads from functional genomics data and the coffee cup sequences and (ii) imputing the genotypes obtained from functional genomics data and the coffee cup sequences at different subsampling rates. The noise levels in the genotypes obtained from subsampled reads and from imputation are depicted by precision and sensitivity plots using the genotypes obtained from WGS of the blood tissue as the true genotypes for RNA-Seq data and six coffee cup samples for each individual. We performed a linkage attack on each subsampled dataset 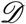 with and without imputation using each subsampled coffee cup sequencing dataset *ℐ* with and without imputation. A total of 2,880 linkage attacks were performed. **(d)** Linking accuracy was calculated for each of the 2,880 linking attacks and depicted as a heatmap. Imputation of the genotypes from functional genomics data improves linking ability, while imputation of the genotypes from coffee cups has the opposite effect. **(e)***gap* values for two individuals and 12 coffee cups each at different sequencing coverage and comparison with *gap* values obtained using random sets of genotypes. The RNA-Seq data used in this plot corresponds to a 20% sampling rate (second row in panel d). This is because the total RNA-Seq data we generated from two consented individuals have deeper sequencing than the gEUVADIS cohort. Twenty percent of the total reads from the study individuals correspond to the total coverage of each RNA-Seq data point in the gEUVADIS cohort, which was selected for fair comparison. **(f)** Cumulative linking accuracy after subsampling the sequencing coverage calculated using the information in panel e. Linking accuracy was calculated with *p*-value cut-offs < 0.01 and < 0.05.

We performed Illumina low-coverage sequencing at 10x coverage for coffee cup samples, which resulted in an average 78.18% and 80.36% of the DNA mapping to the human reference genome for individual A and B, respectively (see Table S1). We called an average of 216,596 and 186,721 genotypes for individuals A and B, respectively. Among them, 55% and 49% of single-nucleotide polymorphisms (SNPs) and small insertions and deletions (indels) in the sample belonged to the individuals A and B, respectively; we calculatedthese percentages using PCR-free WGS data obtained using the blood tissue of the same individuals as the gold standard (see Table S2). We successfully linked all 12 coffee-cup samples to the correct individuals in the dataset with an average *gap* of 1.82 and 2.70 for individuals A and B, respectively, with *p*-values < 10^−2^ (Figure 3b).

#### Effect of imputation and sequencing depth

As mentioned in the previous section, the sequencing depth of RNA-Seq data can vary. However, genotypes missing due to low coverage can be imputed using population genetics-based genotype imputation tools. To understand the effect of sequencing coverage on the noise levels of the attacked database and on the linking accuracy, we subsampled reads from the RNA-Seq data of two study individuals and created cohort genotype datasets *S^D^* for each level of subsampling by using 10, 20,…, 100% of the reads in each RNA-Seq experiment. In addition, we imputed genotypes for each subsampling using BEAGLE [26]. This process resulted in ten datasets *S^D^* from the genotypes of subsampled RNA-Seq reads and ten datasets *S^D^* from the imputed genotypes of subsampled RNA-Seq reads. Similarly, sequencing coverage and the imputation process can greatly affect the noise levels of the query genotypes and, therefore, the linking accuracy. To capture these effects, we subsampled the WGS coverage of coffee-cup samples from 10x to 0.125x and performed genotype imputation for each subsampling. In total, we had six query sets 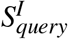 from the genotypes of subsampled WGS reads and six query sets 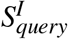 from the imputed genotypes of subsampled WGS reads for each coffee cup. In total, we linked 144 queries (2 individuals x 6 coffee cups x 12 subsample-imputation combinations) to 20 different databases, totalling 2,880 different linkage attacks (Figure 3c).

We first c alculated t he e ffect o f s equencing c overage a nd i mputation o n t he n oise l evels o f the attacked database 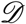 and information *ℐ*. Overall, both the precision and sensitivity of RNA-Seq data increased with the number of reads. However, as reported in the previous section, even when we use all of the reads, the sensitivity and precision of the genotypes from the RNA-Seq data was low. Note that the RNA-Seq data we generated is at a higher sequencing coverage than those in the gEUVADIS cohort. Therefore, the sensitivities of the genotypes in our cohort are higher than those in the gEUVADIS cohort. We found that 20% of the total reads in our experiment was equivalent to the total number of reads of a sample in the gEUVADIS cohort. Additionally, genotype imputation increased the sensitivity of the genotypes called from RNA-Seq by 1,000-fold, but did not affect the precision (Figure 3c). We observed a similar trend when calculating the sensitivity and precision of the genotypes obtained from coffee cups under different sequencing coverages, with and without genotype imputation. Imputation boosted the sensitivity of the genotypes obtained from coffee cups greatly. By contrast, imputation reduced the precision of the genotypes obtained from coffee cups, likely because coffee-cup samples contain multiple individuals’ DNA, which increases the false positive genotypes. This effect is amplified by genotype imputation (Figure 3d).

Next, we assessed how the varying degrees of noise both in the attacked dataset 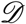 and the query information *ℐ* affect the overall linking accuracy. For each coffee-cup sample (6 cups x 2 individuals), we calculated the linking accuracy for all subsampling and imputation combinations (240 total). We calculated the linking accuracy as the ratio between the number of correctly linked coffee cups and the total number of coffee cups. Figure 3e summarizes the linking accuracy for all scenarios. Overall, we found that imputation of genotypes from the RNA-Seq data improved linking, whereas imputation of genotypes from the coffee cups decreased the linking accuracy. This was especially the case when sequencing coverage was low for both RNA-Seq and WGS. As mentioned above, this result is likely due to the increasing number of false-positive genotypes present on coffee cup samples.

#### Depth-dependent sequencing cost

We investigated the total monetary cost for an adversary to sequence DNA from the coffee cups in order to perform a successful linking attack on a moderately sequenced RNA-Seq cohort. We found that the majority of the coffee cups could be linked with statistical significance to the correct RNA-Seq samples when we spent as little as $19 on sequencing (Figure 3f), with a linking accuracy of 60% (Figure 3g).

#### Different genotyping techniques

An ideal and cheap alternative genotyping method for the adversary in this scenario could be exome-based genotyping arrays, as RNA-Seq captures reads overwhelmingly from exons. To test the results of linking attacks with these methods, we used Illumina exome-based genotyping arrays on the same coffee-cup samples. We found that genotyping arrays had a relatively low call rate for our samples, likely due to fragmented, degraded, and damaged DNA. On average, the call rate was 81% for both individuals (Table S2). Although there were many correctly genotyped SNPs in the coffee-cup samples, their overlap with RNA-Seq genotypes was low, resulting in only 2 out of 12 samples successfully linked to the phenotype dataset (*gap*=1.98 and 1.73, *p*-value< 10^−2^). This result is likely due to the high number of homozygous reference alleles called from the coffee cups (genotype = 0) when we used genotyping arrays. It is difficult to link homozygous reference alleles to RNA-Seq data due to uneven depth distribution and noise. Another genotyping method popular for its low cost and portability is the Oxford Nanopore. SNP and indel genotype quality is quite low with single-pass sequencing using standard protocols, however no study to date has investigated the potential of this sequencing method to be used to violate privacy. DNA extracted from coffee cups is likely partially degraded, highly fragmented, and damaged. We used the simplest and most standard sequencing kit recommended by the Oxford Nanopore with multiplexing (due to the low amounts of input DNA) without the additional steps of DNA repair or suggested quality control (see Methods). The intent of this approach was to minimize the cost and to mimic the act of a curious scientist surreptitiously gathering DNA. On average, we called 65 and 45 SNPs and indels for individuals A and B, respectively. Among these, four and five SNPs and indels were correct for individuals A and B, respectively, but only one SNP was present in the RNA-Seq genotype data (Table S2). Moreover, only a few variants were present in the 1,000 Genomes panel, suggesting that these were false calls. Therefore, linking the coffee cups to RNA-Seq data using a standard Nanopore kit without DNA quality control or damage repair was not successful.

### 2.4 Different functional genomics techniques with varying sequencing depth can be reliably linked to genome databases

Here, we assessed whether linkage attacks yield similar results when data from different functional genomics techniques (e.g., ChIP-Seq, Hi-C, or ChIA-PET) are used as the noisy information *ℐ* and linked to a genome database. Our goal was to empirically quantify and compare the amount of sensitive information in various functional genomics datasets. This is particularly important as different assays target different regions of the genome with different coverage profiles. For example, RNA-Seq targets expressed exons, whereas H3K27ac ChIP-Seq targets the non-coding genome of the promoter and enhancer regions. Different assays also have different coverage profiles (i.e., some assays have spread-out peaks while others are more punctate). We used cell line functional genomics data from the ENCODE consortium as information *ℐ* about individuals represented in the 1,000 Genomes genotype database (the attacked database 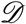). We treated the genotypes in the attacked database 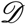 as the gold standard when comparing different metrics such as linking score and *gap* values.

The ENCODE data portal contains data from a range of functional genomics assays (Table S4) on the GM12878 cell line (individual NA12878 in the 1,000 Genomes database). We performed linkage attacks at varying sequencing coverage using data from a number of these assays. We compared our results from functional genomics data to those from WGS, whole exome sequencing (WES), and genotyping arrays. We were able to successfully link individual NA12878 to the 1,000 Genomes database (attacked database 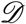) even at very low coverage with Hi-C, total RNA-Seq, polyA RNA-Seq, scRNA-Seq, and ChIP-Seq TF binding data used as information *ℐ*. Overall, ChIP-Seq for histone modification data showed lower gap values at low coverage compared to other assays (Figure 4a). We also calculated the linking score per nucleotide by normalizing the linking scores with the total number of base pairs in an assay. This approach revealed that some of the TF data from ChIP-Seq have higher linking scores per nucleotide than the WGS data (Figure 4b). Although the total linking score obtained using scRNA-Seq data was lower compared to other assays, the *gap* values were surprisingly high even at the lowest coverage (Figure 4a). In general, experiments targeting exons (WES, RNA-Seq) demonstrated comparable *gap* values to whole-genome approaches even though the linking scores were lower. To investigate the reasons behind this, we calculated the median frequency of genotypes called from different functional genomics assays at different coverage (Figure 4c). We found that genotypes from assays targeting exons (especially scRNA-Seq) were slightly more rare in the cohort than genotypes from other assays (Hi-C, WGS, ChIP-Seq TF binding and histone modification), and hence the comparable *gap* values despite lower linking scores (Figure 4c, see Methods for contributions of rare and common genotypes to linking scores). The *gap* value of the polyA RNA-Seq sample to the correct individual at low coverage is higher than the gap value obtained from WES and total RNA-Seq at the same coverage. We think the reason is as follows: polyA RNA-Seq sequences contain highly expressed exons, WES sequences contain all exons, and total RNA-Seq data contain sequences from non-exonic parts of the genes. Compared to all the exons on a gene or other intragenic regions, highly expressed exons contain more rare variants due to selection pressure. Therefore, polyA RNA-Seq data contain more individual specific variants. Thus, although polyA RNA-Seq BAM files typically contain fewer reads than total RNA-Seq or WES BAM files, they can be highly prone to re-identification attacks, which is not clear at first glance.

**Figure 4:**
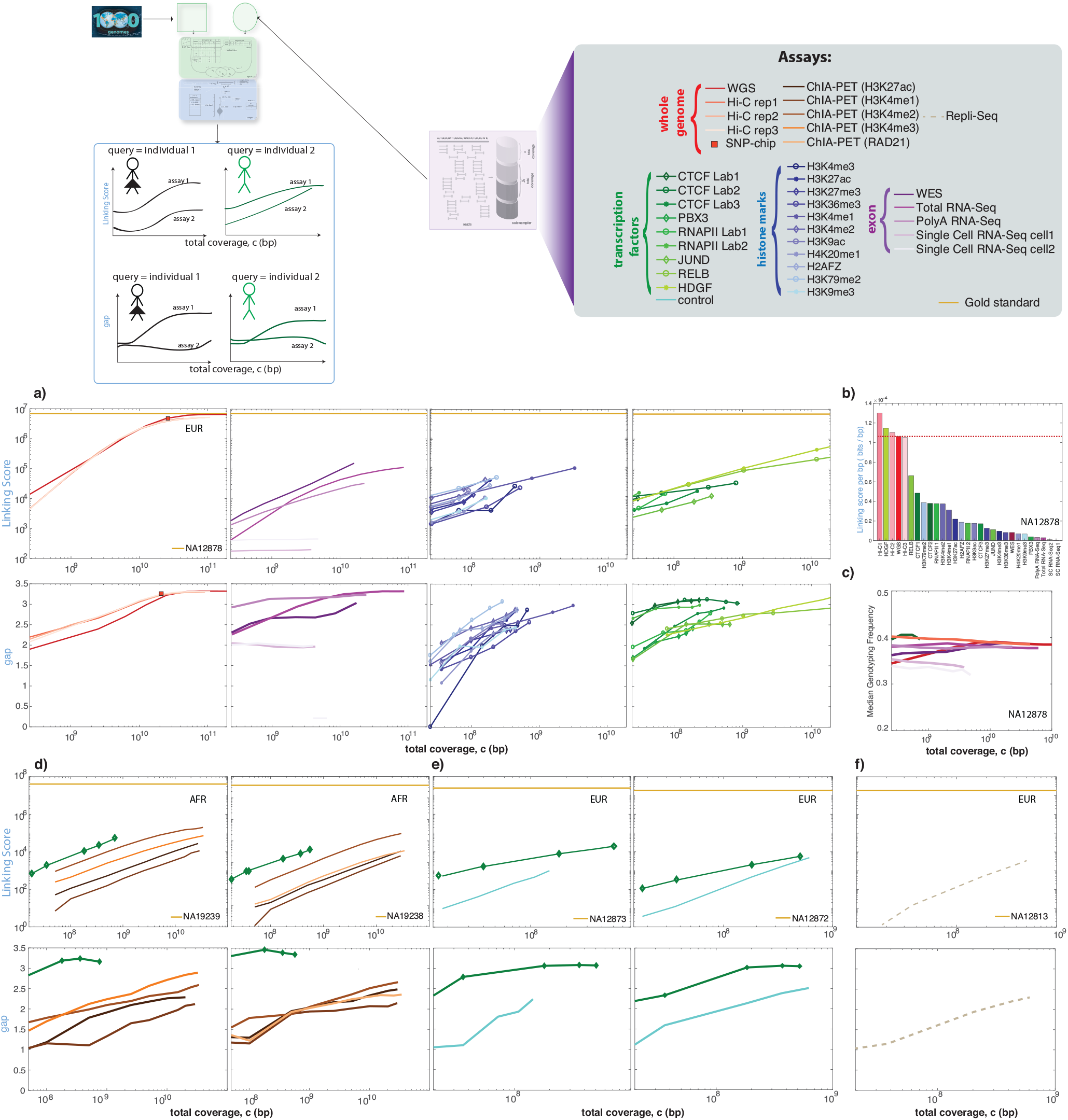
**(a)** The linking scores and *gap* as a function of sequencing coverage for individual NA12878. Different types of functional genomics data were obtained from the ENCODE data portal. **(b)** Linking score per base pair for each assay was calculated by normalizing the linking score values per subsampling by the total number of nucleotides in the subsample. **(c)** Mean genotyping frequency per sequencing coverage for different assays. **(d)** The linking score and *gap* values for ChIA-PET and ChIP-Seq experiments from individuals NA19239 and NA19238 with the 1,000 Genomes genotypes as the gold standard. **(e)** The linking score and *gap* values for ChIP-Seq experiments from individuals NA12873 and NA12872 with the 1,000 Genomes genotypes as the gold standard. **(f)** The linking score and *gap* values for a repli-Seq experiment from individual NA12813 with the 1,000 Genomes genotypes as the gold standard.

We repeated this analysis for individuals for whom we had less functional genomics data. We found that ChIP-Seq experiments targeting the CTCF had high *gap* values even at very low coverage. This held true for every individual in our cohort regardless of their ancestry (Figure 4d-4f). We found that non-obvious data types such as ChIP-Seq control experiments and Repli-Seq data can be used for linking purposes above coverage of around 10 million bp; if we assume a typical experiment has on average a 100 bp read length, then this would correspond to roughly 100,000 reads. Surprisingly, for some of the ChIP-Seq TF binding experiments (information *ℐ*) it was not possible to link the individuals to the 1,000 Genomes database (attacked database 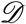) despite their relatively high depth (Figure S2).

#### Effect of ancestral composition of the dataset

We then considered a scenario where we had a panel with a completely different genotype frequency distribution than the 1,000 Genomes panel. To create such a panel, we used genotypes from an African (AFR) population (108 individuals) and added two European (EUR) individuals including NA12878. In this scenario, the *gap* values were still statistically significant and the individual was still vulnerable to linking (Figure S3). We then examined how linking would be affected by removing NA12878 from the panel and leaving in the 108 AFR individuals and 1 remaining EUR individual. Because the genotype frequency of the AFR population is vastly different from the EUR population, we misidentified the remaining EUR individual as NA12878 with a statistically significant gap value (*gap* ∼ 1.6, Figure S3). Our results show that when we have a single individual with the same ancestry as the query individual in a panel, while the rest of the panel has a different ancestry, mispredictions are possible with noisy genotype-based linking. However, the existence of such an imbalanced panel is unlikely.

### 2.5 Practical data sanitization techniques can reduce the amount of private information leakage while preserving the utility of functional genomics data

#### Overview

Sharing read alignment files (SAM/BAM/CRAM) from functional genomics experiments is extremely important for developing analysis methods and discovering novel mechanisms operating on the human genome. Ideally, one would share the maximal amount of information with minimal utility loss while largely maintaining an individual’s privacy. To do so, one must balance the efficiency and effectiveness of the data anonymization process with the utility of the anonymized dataset. Thus, we propose a versatile data sanitization approach such that privacy and utility can be tuned (Figure 5a).

**Figure 5:**
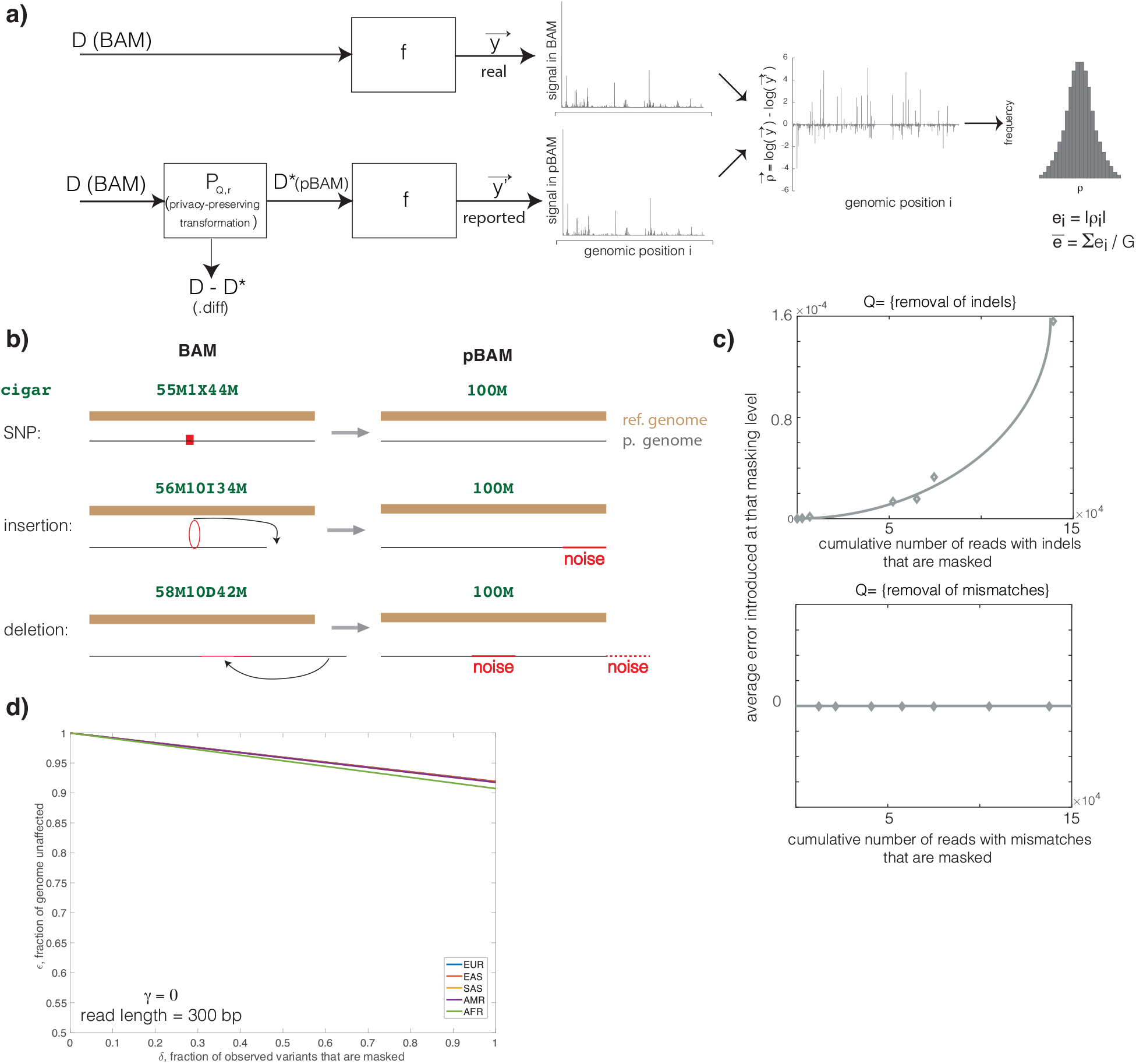
**(a)** The schematic of the privacy-preserving transformation of an alignment file and the difference between the signal calculated from the original BAM and transformed pBAM files. **(b)** A schematic of how different reads and corresponding CIGAR strings are treated in pBAM files. **(c)** The change in error with an increasing number of manipulated reads for different operations *Q*. When *Q* is the removal of mismatches, no noise is added to the depth signal. However, when *Q* is the removal of indels, the error increases with an increasing number of manipulated reads. **(d)** The numerical bounds for the privacy-utility relationship. In the extreme case of obtaining all variants from a functional genomics dataset in an assay with a 150 bp read length, the maximum utility loss is < 10%. The total number of variants and variant types (SNP and indels) were calculated separately for each ancestry represented in the 1,000 Genomes dataset.

A raw alignment file (BAM) can be thought of as a dataset that stores information for each read. Let us assume a BAM file is a dataset *D*, where each entry is a read. The desire is to release dataset *D* in the form of a privacy-preserving BAM (pBAM, say dataset *D*^*^) such that it does not leak variants from the reads, but for which any calculation *f* based on *D* and *D*^*^ returns almost the same result. We achieve this by converting BAM files to privacy-preserving BAM (“pBAM”) files. For example, *f* could be a function that calculates gene expression levels from an input BAM file. The aim is to ensure that the expression levels will be similar, whether BAM 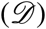 or pBAM (*D*^*^) is used. Now let us perform the privacy-preserving transformation through a function *P_Q,r_* such that *P_Q,r_* (*D*) = *D*^*^. *Q* is an operation such as “removal of variants” and *r* is a parameter representing the number of variants to be removed.

#### Read sanitization protocol

We can construct a sanitized “pBAM” file from a BAM/SAM/CRAM file using generalization (a technique commonly used in data sanitization) for the BAM features. Some of the features in BAM files contain large amounts of variant information that directly reveal the location of a variant (CIGAR and SEQ strings). There are also features that cause subtle leakages (alignment scores, string for mismatching positions, and string for distance to the reference), which allow us to infer the presence of a variant in a read. In general, we address the BAM tags that leak the presence of a variant by generalizing them. For example, we replace the sequence with the corresponding sequence from the reference genome, convert the CIGAR into a format that does not contain variant information, and generalize the other tags. This generalization adds quantifiable noise to the reads depending on the type (mismatches, indels) of the sanitized variants (see Figure 5b, Figure S4, and Methods for quantification of the noise). Our theoretical (Methods) and empirical (Figure 5c) quantifications demonstrated that pBAM adds noise to the number of reads at certain loci when there are indels, and does not alter this number with mismatches. The details of pBAM construction can be found in the Methods and Supplemental Information.

We store removed information in a compressed file format called “.diff” (see Supplemental Information). These.diff files are small files to be kept behind controlled access. To keep the size of private file formats relatively small, we report only the differences between BAM and pBAM in the.diff file and avoid printing any sequence information that can be found in the reference human genome. This allows us to share the majority of the data with minimal utility loss. If the users find the data useful for their research, they can then apply to access the information in the“.diff” files. We provide an easy-to-use software suite that can successfully convert pBAM files to their original BAM format.

#### pBAM provides privacy while maintaining utility

Let us assume that the number of variants to remove is *r*^t^. Our sanitizer *P_Q,r_* will in fact remove *r* number of variants (*r* ≥ *r*^t^), because the variants in linkage disequilibrium (LD) with the *r*^t^ variants need to be removed as well (as one could impute variants within LD blocks). However, note that if the goal is to sanitize all variants from the BAM file, then the LD is irrelevant since all the observable variants will be removed from the BAM file.

We define key quantities to measure privacy and utility. Here, we define privacy of a pBAM with a parameter *δ* and consider the resulting pBAM to be *δ*-private. *δ* is the proportion of sanitized variants to the total number of observable variants (*δ* = 1 means 100% privacy, i.e., all the observable variants are sanitized; see Methods for details). We define the utility of a pBAM with another parameter *ε* and consider the resulting pBAM to have *ε*-utility. *ε* is the proportion of unchanged genomic units to the total number of units based on an error per unit (*e_i_*) threshold *γ*. Error per unit (*e_i_*) is the log-fold difference between the value of the unit in the pBAM vs. BAM formats (see Methods for formulation). For example, if we calculate the signal depth profile from a BAM and a corresponding pBAM, some of the bases in the genome (i.e., units) will have different values. This difference is quantified as added error (*e_i_*, Figure 6a, Methods). If this error is above a threshold value (*γ*), then a given base will be considered as changed. The total proportion of unchanged bases to the length of the genome will then be the *ε* value for this pBAM. Error and utility can also be defined at the level of genes or functional elements. The difference between a BAM and pBAM file at nucleotide resolution gives us the upper bound for the utility. *ε* = 1 means 100% utility, i.e., the results obtained from a BAM and a PBAM are identical.

**Figure 6:**
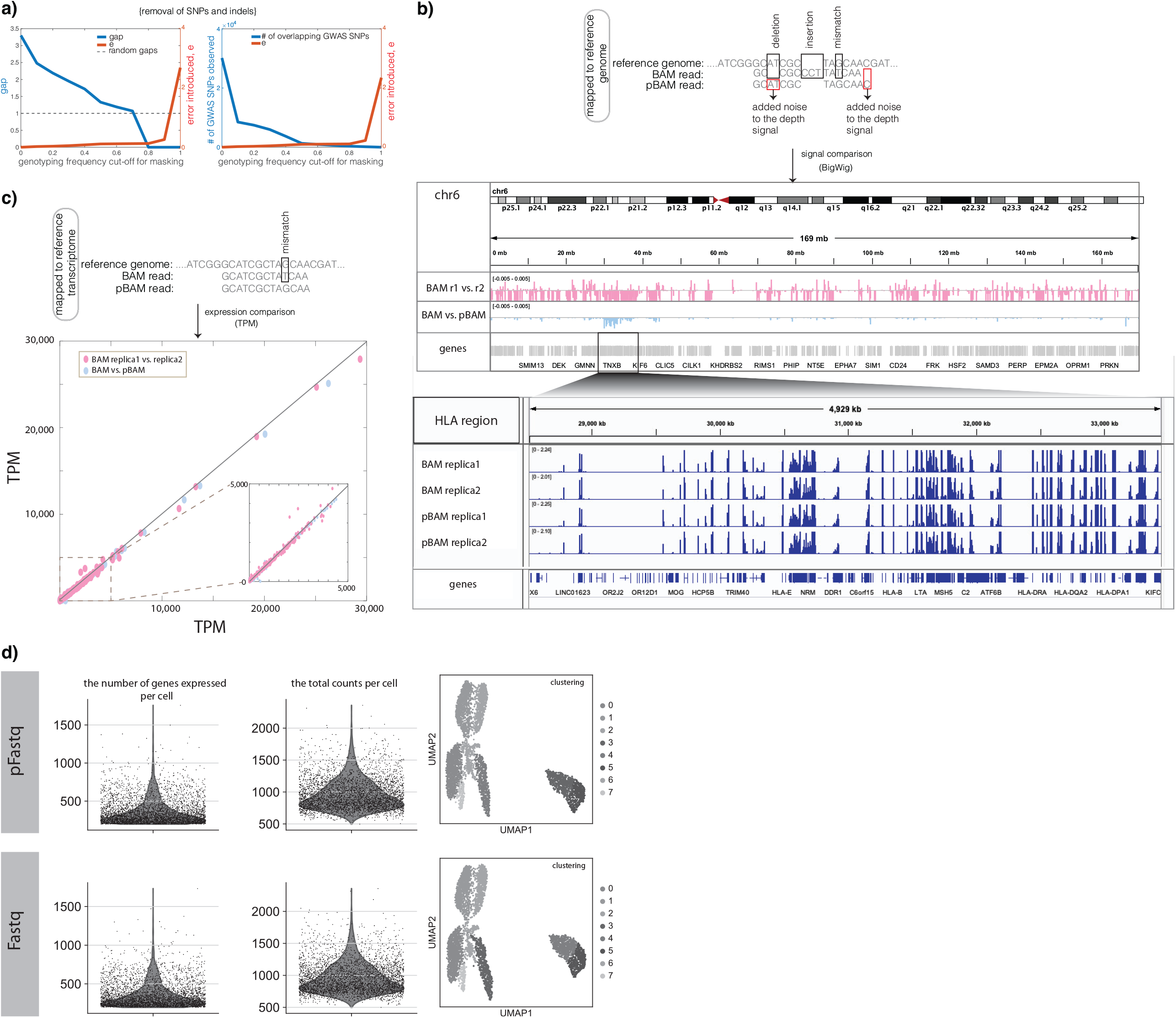
**(a)** The empirical values for privacy-utility balance using NA12878 total RNA-Seq BAM files. We removed the variants with increasing genotyping frequency from the BAM files and calculated the *gap* values and total error in the signal profiles for each pBAM. We also calculated the number of overlapping GWAS variants vs. the total error of the resulting pBAM files. **(b)** The difference between the BAM and pBAM files is shown at the read level, when the reads were mapped to a reference genome. The added noise to the depth signals due to the BAM-to-pBAM transformation is shown in an example read with a deletion, insertion, and mismatch. The difference of the depth signal when calculated from BAM and pBAM is shown in the signal tracks for chr 6. Log-fold differences of the signal of each nucleotide between the BAM vs. pBAM and BAM of replica 1 vs. replica 2 were calculated using deeptools [37]. **(c)** The difference between the BAM and pBAM files is shown at the read level, when the reads were mapped to a reference transcriptome. The BAM files from two replicas and pBAM were then used to quantify the expression of all 60,699 genes. Transcripts per million (TPM) values were compared between the replicas and between BAM and pBAM. **(d)** The difference between the BAM and pBAM files is shown when the aligned reads were converted back to fastq files for 10x v2 single-cell RNA-Seq data. Fastq and pFAStq files were processed using the HCA’s optimus pipeline. The comparison between fastq and pfastq processing is shown for the number of genes, total number of counts, and cell clustering levels.

We derived a mathematical relationship between the privacy parameter *δ* and the utility parameter *ε* to clarify the trade-off between the privacy and utility of a pBAM (see Methods). This relationship depends on the type of sanitized variant, the number of sanitized variants, and the read length. Hence, the privacy-utility relationship will be different for different BAM files as well as for individuals with different ancestries. Therefore, to give one sense of the trade-off, we empirically calculated this relationship for the upper bound as follows: We first assumed that all of the genotypes of an individual can be observed by any functional genomics data. This is a generous assumption as, for instance, one can call around 0.5% of all the genotypes from a typical RNA-Seq BAM file. We then calculated the mean number of SNPs and indels of all individuals from the same ancestry using the 1,000 Genomes database. Finally, we calculated the change in utility as a function of the change in privacy (unit = nucleotide, *γ*=0) (Figure 5d). We found that, at 100% privacy, the utility loss is at most less than 10% for all the ancestries (Figure 5d). We found a small difference in utility loss (∼2%) when we used the number of variants in the African population. This can be explained by the large difference in the number of variants in the African population compared to other populations (Figure 5d).

One way to understand how to interpret the values of the key utility quantities–*e*, *γ*, and *ε*–is to compare them against the discrepancies between the replicates. It is well known that high-throughput experiments such as functional genomics assays are subject to great variability. This is remedied by using biological replicates in experiments and further performing irreproducibility discovery rate analysis [27]. One can calculate the *e_i_* values for the discrepancy between the replicates and use them as the *γ* threshold, i.e., the tolerable error between BAM and pBAM. In particular, if the difference between replicates per unit is 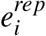 a pBAM can be considered to have no utility loss up to this 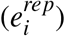 ntity (see Methods).

#### Empirical calculations validate privacy provided by pBAM

To test the assumption that variants are masked in pBAM files, we performed variant calling of the pBAMs at different privacy levels on an RNA-Seq data of individual NA12878. We first systematically ramped up the total number of masked variants by thresholding using the genotyping frequencies in the 1,000 Genomes database. We then performed variant calling and subsequent linkage attacks on the database and calculated the *gap* values. We also calculated the number of overlapping GWAS SNPs with the called variants from pBAM set as another privacy metric. Figure 6a shows that variant calling yields a lower number of genotypes, hence a lower number of overlapping GWAS SNPs and fewer *gap* values with pBAM at different privacy levels.

#### Empirical calculations validate utility provided by pBAM

We further calculated the total amount of error introduced in the signal depth profiles when we convert the BAM file to a pBAM file at each genotyping frequency thresholding level. Figure 6a shows how risk of privacy loss decreases with increasing loss of utility. To further understand utility loss, we calculated the difference (log2 ratio) between the signal depth profiles (BigWig files) obtained from pBAM and BAM files when we masked all of the variants. In addition, we performed the same comparison between the signal depth profiles obtained from BAM files belonging to two different biological replicates. As shown in Figure 6b, the difference between BAM and pBAM files is smaller than the difference between the replicates at base resolution, suggesting that the noise added to the signal is within the biological noise levels. We zoomed in to the highly polymorphic HLA locus to show the similarity between the signal in biological replicates and their corresponding pBAMs. We further quantified the gene expression levels and found minimal difference between pBAM and BAM files (Figure 6c). We performed similar calculations for ChIP-Seq data (Figure S5).

#### Implementation

We implemented our pipeline for converting between BAM and pBAM+.diff files (Figure 7a) in bash, awk, and Python. The.diff files are encoded in a compressed format to save disk space. For convenience, pBAM files are saved as BAM files with manipulated content and with a p.bam extension. That is, any pipeline that uses BAM as an input can take p.bam as an input as well. CPU times (calculated using a single 2.3 GHz AMD Opteron processor) and associated file sizes for alignments from RNA-Seq and ChIP-Seq experiments are shown in Table 1. Our data sanitization pipeline has been adopted by the ENCODE Consortium Data Coordination Center and deployed in the ENCODE Uniform Pipeline Framework using workflow description language scripts and docker images, accompanied by appropriate documentation for computational reproducibility on multiple platforms (Google Cloud, Slurm Scheduler, LINUX servers, etc.) under ENCODE Data Processing pipelines. Codes for calculating information leakage, scripts for file manipulations, examples, and file specifications of BAM, pBAM, pCRAM, and.diff files can be found at privaseq3.gersteinlab.org and github.com/ENCODE-DCC/ptools.

**Table 1:** p-tools performance and associated file sizes

| **Experiment** | Total RNA-Seq | PolyA RNA-Seq | ChIP-Seq (CTCF) | ChIP-Seq (H3K4me1) |
| --- | --- | --- | --- | --- |
| **BAM size (bytes)** | 35,219,346,385 | 16,301,017,652 | 954,993,667 | 2,230,202,265 |
| **pBAM size (bytes)** | 31,986,293,946 | 14,057,962,755 | 876,237,603 | 1,838,044,304 |
| **CRAM size (bytes)** | 21,317,709,703 | 10,084,587,074 | 425,193,485 | 1,120,941,477 |
| **pCRAM size (bytes)** | 20,436,579,905 | 966,7758,517 | 378,736,942 | 883,158,959 |
| **.diff size (bytes)** | 623,943,745 | 260,978,063 | 6,991,519 | 19,438,692 |
| **BAM to .diff compression** | 99.98% | 99.98% | 99.99% | 99.99% |
| **CRAM to .diff compression** | 96.94% | 97.30% | 99.98% | 99.98% |
| **BAM+hg to pBAM+.diff CPU time** | 23:21:17 | 17:05:26 | 00:37:11 | 01:08:00 |
| **pBAM+.diff+hg to BAM CPU time** | 12:34:23 | 04:21:05 | 00:27:47 | 00:46:50 |

**Figure 7:**
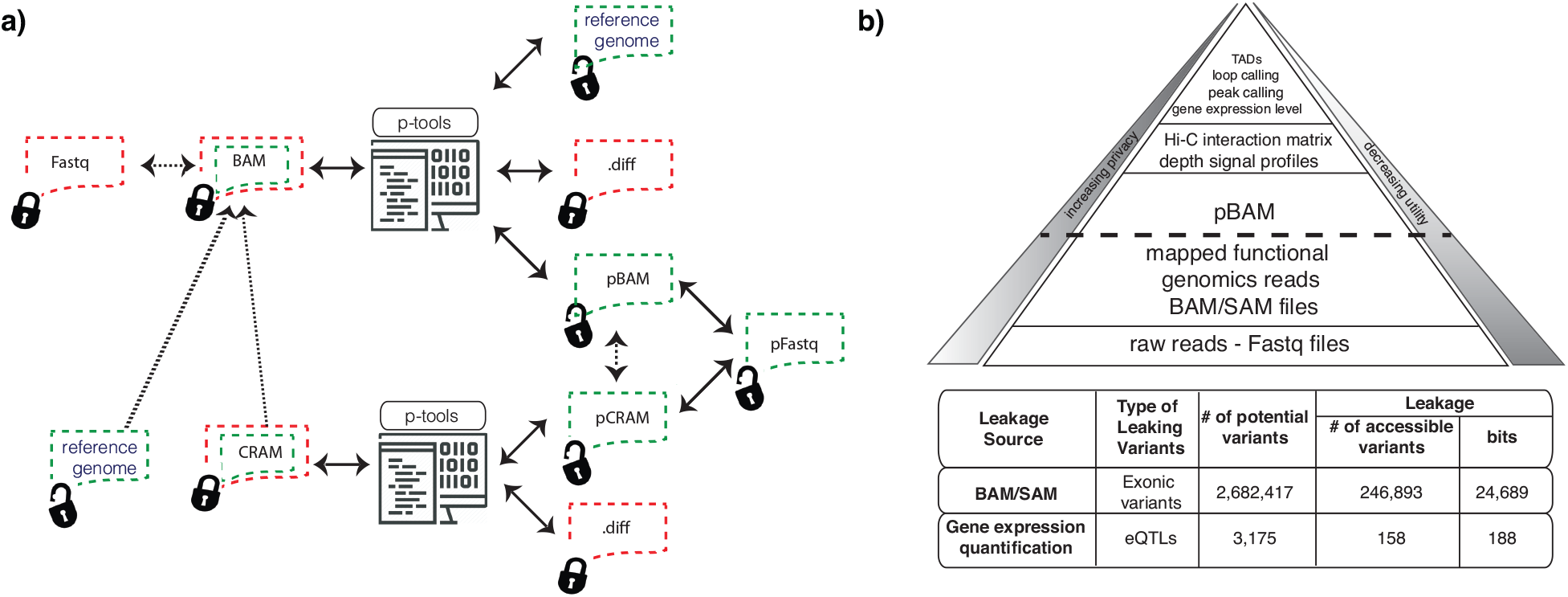
**(a)** Schematic of how p-tools work with different file formats. **(b)** Different layers of data produced from functional genomics experiments. Raw reads from the sequencer that are stored as FASTQ files are mapped to the reference genome. These data types leak the greatest amount of sensitive information, while also possessing the highest utility. Moving upwards through the pyramid levels, there is less privacy concern but largely reduced utility and amount of data.

#### pFastqs as an alternative for alignment-free expression quantification and 10x scRNA-Seq reads

There are many different ways to quantify expression levels using RNA-Seq reads. Some software such as RSEM [28] use aligned reads (BAM files), while others such as kallisto [29] use reads directly from fastq files. While converting pBAM files to pFastqs for bulk functional genomics data is trivial with available tools (e.g., samtools), 10x scRNA-Seq reads have different fastq structures. To address this, we added a module to our software suite that first maps the 10x scRNA-Seq reads to the reference genome and transcriptome, and generates a BAM file that includes the unaligned reads (following a typical 10x scRNA-Seq pipeline), as well as the tags that contain information about barcodes and unique molecular identifiers (UMI). Our software then converts this BAM file into pFastq files based on UMI, barcode, and sequence identifiers. By using a 10x (v2) scRNA-Seq dataset from the Human Cell Atlas project, we calculated the key quantities using fastq and pFastq files as input (Methods). We found that both fastq and pFastq returned similar results in terms of total gene count, total counts per cell, and cell clustering. Note that clustering algorithms are often used stochasticity and may return slightly different results even when the same input is used (Figure 6d).

## 3 Discussion

Functional genomics experiments are increasingly leveraging tissues or cells from donors due to their clinical significance. For example, large-scale studies such as TCGA, GTEx, HCA, and PsychENCODE provide functional genomics data on thousands of human subjects with known phenotypes such as cancer or psychiatric patients. We live in the era of the “omics revolution” and expect to have a surge of more and more functional genomics data on human subjects in the near future, which will require data access solutions beyond the traditional approaches in place right now. Here, we aimed to provide privacy- and utility-preserving solutions to the functional genomics data access problem faced by the field.

As with all experimental data, sharing “raw” functional genomics data is important because it allows researchers to obtain biological insights by leveraging their own tools and analyses. The availability of raw data also helps to overcome the reproducibility crisis in scientific research. However, the majority of functional genomics experiments use next-generation sequencing-based assays; hence, their raw form involves pieces of sequences from donors’ genomes. In order to preserve patients’ privacy while providing high-utility data, the field urgently needs solutions to data access.

We designed a data sanitization protocol that converts functional genomics BAM files into pBAM files, which masks the variants in the reads while maintaining the utility of these files. Our data sanitization protocol is flexible and based on principled trade-offs between privacy and utility. While the most privacy-conservative option (i.e., 100% private) is to remove all split reads (to avoid inference of structural variants) and convert all SNPs and indels to the reference, the tool allows study participants to choose to be more or less conservative. For example, some participants may wish to mask only the variants that leak information about their susceptibility to stigmatizing phenotypes or diseases that can be used against them by insurers or employers. In that case, researchers can convert BAM files to pBAM files by masking only the desired variants and those in LD, and theoretically calculate the utility loss. To help guide study participants and researchers, we developed a formalism to show information utility loss under different coverages along the genome. For example, the genomic coordinates of a non-expressed gene will not be sequenced in RNA-Seq, so removing any indel overlapping with these locations will not result in any utility loss. Our formalism will allow participants and researchers to find an optimal combination of variants to be masked in order to maintain high utility. For the remaining unmasked variants, our linkage attack software can be used to assess the empirical privacy risks of sharing.

To design effective data-sanitization protocols, one needs to quantify the private information leakage in the data. One way to achieve this is through linkage attacks, which provide means to quantify the private information leakage in a database. For example, as was shown previously, anonymized identifiers are not enough to anonymize DNA sequencing data, as the information in DNA can be cross-referenced with external databases to infer potentially private information about the participants [6].

Deriving important quantities such as gene expression values or TF binding site enrichment does not require the knowledge of the genetic variants of a sample. Our aim was to understand how we could mask the genetic variants in the raw functional genomics reads without perturbing the resulting analysis. To achieve this, we first needed to systematically quantify genetic variant leakage in raw functional genomics data and analyze the robustness of the leakage under different circumstances. We demonstrated how potentially private phenotypes can be inferred by de-anonymizing raw functional genomics data through instantiating linking attacks using DNA information from known individuals. This effort also revealed an important and unique aspect of DNA linkage to the functional genome: the coupling between the data and phenotype associated with a given sample. We also illustrated how DNA is becoming more and more accessible to the average “citizen scientist”; for just $19, we were able to link genotypes obtained from sequencing a consented individual’s used coffee cup to their anonymized raw RNA-Seq reads.

We addressed the most obvious leakage from functional genomics data, and provided solutions for quick quantification and safe data sharing. Other sources of information leakage from functional genomics experiments are possible at different stages of the data summarization process (Figure 7b). Subtle leakages can come from the quantification of expression values; given a population of individuals, these gene expression values can be related to variants through eQTLs, and hence can create leakage [12, 11, 13]. In Figure 7b, we calculated the potential number of variants one can obtain (i.e., the total leakage) from a typical RNA-Seq experiment in order to get a sense of the contribution each type of leakage makes to the total leakage (details can be found in Figure S6). While inferring the leakage from gene expression values through eQTLs is interesting and non-trivial, eQTLs are not a main source of genotype information in functional genomics data. We showed that the amount of genotype leakage from raw alignment files is almost ∼ 1, 000 times that from gene expression levels, and can be avoided with pBAMs. Another source of subtle private information leakage could be the presence of viral (e.g., HPV/EBV) or bacterial reads in raw alignment files. Future studies could focus on the quantification of privacy loss from functional genomics data. Nevertheless, inference of potentially stigmatizing phenotypes can be avoided by simply removing these reads from the BAM files during the data sanitization process.

## 4 Methods

### 4.1 Linkage Attack Details

#### 4.1.1 Linking score

Let us assume that there are *n* variants that can be observed from our query individual with known identity, either by using deeply sequenced WGS data (perfect information *ℐ*) or by using DNA left on a coffee cup (noisy information *ℐ*). 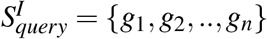 is then the set of genotypes for each variant (*g_i_* = {0, 1, 2} for the homozygous reference allele, heterozygous alternative allele, and homozygous alternative allele, respectively). 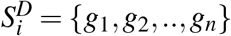 is the set of genotypes for an individual *ℐ* in dataset 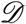 for the same variants in 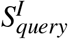. Note the genotype set 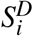 is observed from functional genomics data of an anonymized individual *ℐ* and some of the genotypes may be missing due to the coverage and experimental bias in the functional genomics reads. For each individual *i* in *S^D^*, we find the intersection 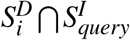and calculate a linking score

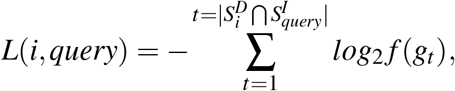

where *f* (*g_t_*) is the ratio of the number of individuals whose *t^th^* variant has the genotype *g_t_* to the total number of individuals in the cohort.

#### 4.1.2 gap

To find the individual in dataset 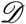 that matches the query individual, we then rank all the *L* (*i, query*) scores for a given query individual in decreasing order as

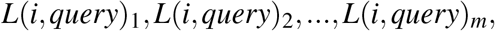

where *m* is the total number of individuals in the attacked dataset. We denote the individual with the highest score as the queried individual. To assess the statistical robustness of this prediction, we defined a measure called *gap*, which is the ratio between the *L* (*i, query*) score of the first-ranked individual (*max* = *L* (*i, query*)_1_) and that of second-ranked individual (*max*_2_ = *L* (*i, query*)_2_ and *gap* = *max/max*_2_). The goal is to determine how separated the matching individual is from the rest of the individuals in the cohort. The *gap* value is similar to the eccentricity parameter in the IMDB-Netflix attack [14] used to determine the reliability of the linkage attack. The difference between *gap* and eccentricity is that *gap* is the fold change, which can be compared across samples.

##### Statistical significance of gap

We estimated the empirical *p*-values for a particular *gap* value by linking a set of random genotypes that do not belong to a particular individual to the database, as follows.

a. We selected *n* random genotypes from a database of nearly 50 million genotypes (taken from 2,504 individuals in the 1,000 Genomes database). *n* is the total number of genotypes that were linked to the database for the *gap* calculation.
b. We calculated the *L* (*i, query*) between these random *n* genotypes and every individual *i* in the database.
c. We calculated the gap as the ratio between the first-ranked and second-ranked *L* (*i, query*).
d. We repeated the above steps 1,000 times and obtained a distribution of random *gap* values.
e. The total number of random *gap* values that are equal to or greater than the real gap divided by 1,000 is the probability of observing the real *gap* value by chance.

We empirically calculated the probability of observing the real *gap* value or a larger *gap* value by chance by randomly subsetting *n* variants (the same number of variants as in 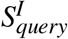) observed from the information *ℐ*, performing the linkage attack, and calculating the associated *gap* value. If this *p*-value is statistically significant, then the attacker can rely on the prediction.

#### 4.1.3 Addition of false-positive genotypes to the attacked database

Using the 1,000 Genomes dataset, we randomly selected genotypes and appended them to the genotype set of each individual *i* 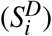. These random genotypes may or may not be found in the genotype set of 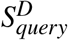, hence increasing the chance of matching the query individual to any individual *i* in *S^D^*.

### 4.2 Sample Selection

We present short variant calls on 478 samples. Of these, 16 were newly sequenced for this study (two DNA samples from two individuals, two RNA samples from the same individuals, six coffee cup DNA samples from two individuals), and the remaining 462 RNA-Seq data were obtained from the gEUVADIS study [23]. Genomic materials for newly sequenced samples were obtained by collecting blood samples and used coffee cups. DNA samples from the blood were sequenced using high-coverage Illumina sequencing (30x) and used as the gold standard. RNA samples from the blood were sequenced using the Illumina total RNA-Seq protocol. Extracted DNA from coffee cups were sequenced using low-coverage Illumina sequencing (10x), Oxford Nanopore Technologies (ONT), and genotyped with the Illumina Infinium OmniExpressExome-8 v1.6.

### 4.3 Contribution of very rare and unique genotypes to the *L*(*i*, *query*) score

We calculated the number of unique, very rare, and common genotypes for every individual in the 1,000 Genomes panel. We observed around 15,000 unique genotypes per individual. This contributes around 11 × 15, 000 = 165, 000 bits of information. We estimated 11 from −*log*_2_(1/2503), as 1 in ∼2,503 individuals in 1,000 Genomes have these unique genotypes. We observed around 670,000 very rare genotypes, which contribute 7 × 670, 000 = 4, 690, 000 bits of information on average. We estimated 7 from −*log*_2_(20/2503), as 20 in ∼2,503 individuals in 1,000 Genomes have these unique genotypes. In total, unique and very rare genotypes contribute 4, 855, 000 bits of information. We then calculated the information in the genomes of all the individuals in the 1,000 Genomes Phase III panel. The mean information per individual is around 2*x*10^7^ bits. The contribution of unique and very rare variants then becomes around 24% of the total information in an individual’s genome, despite the fact that the number of unique and very rare variants is only 3% of the total number of variants in an individual’s genome. Note that this calculation is based on our scoring system adopted from Narayanan and Shmatikov [14], which assumes independence between variants.

### 4.4 Experimental protocols

#### 4.4.1 DNA extraction protocol from coffee cup lids

We used the QIAamp DNA Investigator Kit from QIAGEN. This kit is designed to purify DNA from forensic and human identity samples. We first swabbed the surface of the coffee cups using a cotton swab dipped in 1 *μ*L purified water. We followed the QIAamp DNA Investigator kit protocol for isolating DNA from surface-swab samples without modification. The final amount of DNA isolated from coffee cups was around 0.9 to 1 ng.

#### 4.4.2 Whole-genome amplification

Due to the very low starting amount of purified DNA, we used a single-cell whole-genome amplification kit (REPLI-g Single Cell Kit), which allows uniform PCR amplification from single cells or limited sample materials for use in next-generation sequencing applications. We then used the Monarch PCR and DNA Cleanup kit to purify the DNA from PCR reactions.

#### 4.4.3 Illumina sequencing

Amplified DNA samples from coffee cups as well as purified PCR-free DNA from blood (as the gold standard) were sent to the Yale Center for Genome Analysis for Illumina WGS. Coffee cup samples were sequenced at a 10x coverage and blood samples were sequenced at a 30x coverage.

#### 4.4.4 Illumina genotyping arrays

We used an Infinium OmniExpressExome-8 BeadChip for the amplified DNA samples from coffee cups. Infinium OmniExpressExome-8 array surveys tag SNPs located on exons from all three HapMap phases, which includes 273,000 exonic markers. Each SNP is represented on these chips by 30 beads, on average. The Yale Center for Genome Analysis performed the BeadChip protocol and calculated the call rates using an Illumina BeadStudio.

#### 4.4.5 Nanopore Sequencing

Due to the low quality of the DNA obtained from coffee cups, we did not perform size selection of the fragments in the PCR-based libraries we obtained using the ONT rapid sequencing kit. A total of 12 libraries from six coffee cups per individual were barcoded using the ONT rapid barcoding kit. Libraries were sequenced across an individual R9.4 flow cell on a single MinION instrument. A total of 844,599 reads were successfully base-called and demultiplexed using Guppy. The recommended MinION run-time was 48 h; therefore, the run was terminated after 48 h. SNP calling was performed using Nanopolish software.

#### 4.4.6 RNA extraction protocol and RNA-Seq

Blood samples from individuals were sent to the Yale Center for Genome Analysis for RNA purification and Illumina high coverage total RNA-Seq analysis following the suggested protocols by Illumina. Total RNA-Seq data yielded more genotypes than the gEUVADIS data. To do a fair comparison for the linkage attacks, we downsampled the total number of captured variants to the average number of variants observed in the gEUVADIS dataset.

### 4.5 Genotyping

#### Blood tissue and coffee cups

##### Illumina sequencing

The DNA extracted from two blood tissues and 12 coffee cup samples were sequenced using Illumina. Raw fastq files were processed by mapping them to the hg19 reference genome (b37 assembly) using bwa [30]. The resulting BAM files were processed using Picard tools to remove PCR duplicates. De-duplicated files were then genotyped using GATK best practices [21, 22].

##### Genotyping Arrays

The 12 coffee cup samples were genotyped using Illumina Infinium OmniExpressExome-8 v1.6 and UV-coated chips were scanned using IScan. Scanned output files were analyzed and call rates were calculated using Illumina BeadStudio.

##### ONT

The 12 coffee cup samples were prepared and sequenced using the rapid barcoding kit and minION following the manufacturer’s suggestions. After converting fast5 files to the fastq format using Guppy software for base-calling and de-multiplexing, each sample was aligned to the hg19 reference genome (b37 assembly) using bwa “mem −x ont2d” options [30]. Aligned reads were used for variant calling using Nanopolish software.

##### Functional genomics data

Raw RNA-Seq fastq files (from this study, gEUVADIS, and ENCODE) were processed by mapping them to the hg19 reference genome (b37 assembly) and gencode v19 transcriptome assembly using STAR [31]. Other raw functional genomics (e.g., Hi-C, ChIP-Seq) fastq files were mapped to the hg19 reference genome (b37 assembly) using bwa [30]. The resulting BAM files were processed using Picard tools to remove PCR duplicates. Deduplicated files were then genotyped using GATK best practices [21, 22] for RNA and DNA for RNA-Seq and other functional genomics data, respectively.

### 4.6 pBAM details

The privacy-preserving BAM (pBAM) format is a variant of the BAM format. The genomic variants in BAM files are masked using data-sanitization techniques as outlined below. pBAMs are generated after aligning the functional genomics reads to the reference genome. The alignment step can be performed in a privacy-preserving manner as well [32], however the resulting alignment files still need to be sanitized to maintain patient privacy. The goal is to create files that can be openly shared and utilized in any downstream analysis pipeline. Our data sanitization is agnostic to the type of variant. Any difference between the sample and the reference genome will be covered regardless of whether they are somatic or germline SNPs or large structural variants. We simply compare the read in a BAM file to the reference genome and remove the difference. This will prevent genotyping and variant calling from the reads [33].

#### 4.6.1 Procedure

Below is a practical guide on how to convert BAM files to pBAM files.

1. Let us assume the variants that need to be sanitized from the BAM file are *V_s_* = {*s*_1_*,‥, s_i_,‥s_n_*} and the variants that are in LD with the variants in *V_s_* are 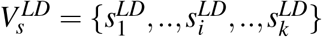. The total number of variants that need to be sanitized are then *r* = *n* + *k*.
2. We first find all the reads that contain the variants in the 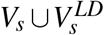 such that *R* = {*R*_1_*,‥, R_T_*}.
3. We apply sanitization techniques (i.e., generalization) to the BAM fields so that an adversary cannot infer the existence of these variants:

a. CIGAR: Convert the CIGAR of each *R_i_* to a perfectly mapped read CIGAR (e.g., 100M, where 100 is the read length and M denotes that each base on the read perfectly mapped to reference genome).
b. SEQ: Replace the sequence of each *R_i_* with the sequence in the reference genome.
c. QUAL: Convert all the base qualities of *R_i_* to perfectly called base phred scores.
d. There are also optional tags in the BAM files such as AS (alignment scores), MD (string for mismatching positions), and NM (edit distance to reference) that should be sanitized. They can be generalized to the values for perfectly mapped reads if they are present in the BAM files.
e. We demonstrated that there might be extremely subtle leakages through MAPQ scores (mapping quality, see Figure S7). In particular, if the goal is to prevent large leakages such as structural variants, then a data-sanitization procedure such as suppression or generalization might be suitable for this field as well.

In addition, we treat intronic reads differently to be able to capture the splicing accurately. Details can be found in the Supplementary Information.

#### 4.6.2 Privacy

If one can observe *t* number of variants from a functional genomics BAM file, then the resulting *D*^*^ = *P_Q,r_*(D) can be viewed as *δ*-private with respect to operation *Q*, if *δ* = *r/t*, which is the ratio of non-observable variants in a pBAM file to all observed variants in a BAM file. As mentioned earlier, *P_Q,r_* is the data sanitizer. We can reach 100% privacy when *r* = *t*. Here, variants are defined as homozygous and heterozygous alternative alleles. This definition assumes that all *r* number of variants are guaranteed to be deleted in the resulting pBAM file.

#### 4.6.3 Utility

We defined the the utility of pBAMs as such that any calculation *f* based on a BAM file should result in similar results when the pBAM is used instead. A calculation *f* can be a signal depth profile calculation, TF binding peak detection, or gene expression quantification (Figure 5a-5b).

Then, we can reconstruct an equation for each unit *i* as

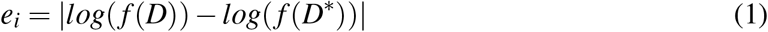

where a unit *i* can be a single base pair, an exon, or a gene depending on the function *f*. Accordingly, *e_i_* can be calculated as the absolute value of the log-fold change between the results derived from the BAM and pBAM file. Note that *e _i_* is a measure of the error in the new dataset *D*^*^. *D*^*^ = *P_Q,r_* (D) can be viewed as having *ε*-utility with respect to operation *Q* if *ε* = (*G* − *m*)*/G*, where *m* is the total number of units with *e_i_ > γ* and *G* is the total number of the units. We can obtain 100% utility if the error is smaller than the threshold *γ* for every unit in the genome.

##### Concordance between replicates vs. BAM and pBAM

In an effort to avoid assigning the *γ* threshold *ad hoc*, we can use the concordance between biological replicates of a functional genomic experiment as guidance. Let us denote *R*_1_ and *R*_2_ as the BAM files from two biological replicates of the same experiment. Then, we can repurpose the equation above as

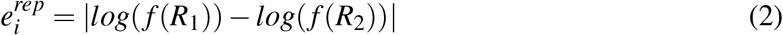

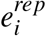 is the absolute value of the log-fold change between the results derived from *R*_1_ and *R*_2_. Then, the *γ* for every unit can be chosen as 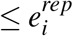, such that the error tolerated between BAM and pBAM never exceeds the biological noise levels.

#### 4.6.4 Privacy-Utility Relationship

The relationship between privacy and utility can be derived through the mathematical relationship between *δ* and *ε*. Let us assume the units for our utility calculation are single bases in the genome, as this will give us the upper bound (i.e., more utility loss at higher resolution). Let us also assume that the function *f* for which we want to measure the utility loss is the signal depth calculation.

Sanitization is done over three kinds of variants: SNPs, insertions, and deletions. (1) In the case of SNPs, when we change a letter from the alternative allele to the reference allele, the resulting signal profile at that location does not change. (2) In the case of an insertion at position *x*, when we delete the insertion from a read (since an insertion is not represented in the reference), we have to append *l_ins_* number of bases to the end of the read (*l_ins_* is the length of the insertion). This adds error to all of the bases between position *x* and *x* + *L_R_*, where *L_R_* is the length of the reads. That is, for each insertion, the total number of bases with *e_i_ > γ* will be at most *L_R_*, when *γ* is equal to 0 for the upper bound. (3) In the case of a deletion at position *x*, when we fill the deletion with the reference, we have to delete *l_del_* number of bases from the end of the read (*l_del_* is the length of the deletion). This adds error to all of the bases between position *x* and *x* + *L_R_* + *l_del_* − 1, where *L_R_* is the length of the reads. The maximum detected indel length varies by the aligner settings. In the most extreme cases, *l_del_* can be as large as *L_R_* − 1. That is, for each deletion, the total number of bases with *e_i_ > γ* will be at most 2 · *L_R_* − 2, when *γ* is equal to 0 and *l_del_* is equal to *L_R_* − 1 for the upper bound.

If *r* is the total number of variants to be sanitized, then *r* = *r_snp_* + *r_ins_* + *r_del_*. The number of bases with *m* such that *e_i_ >* 0 are at most

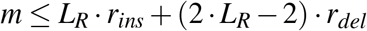

Since *ε* = (*G* − *m*)*/G*, then *m* = −*ε* · *G* + *G*. We can then say

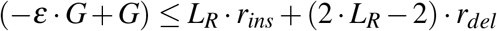

If we replace *r_ins_* with *r* − *r_snp_* − *r_del_* and *r* with *δ* · *t*, then our relationship becomes

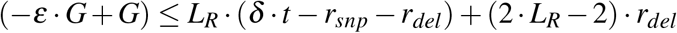

#### 4.6.5 Utility bounds

The goal is to consider the impact of sanitizing BAM files through modifications applied to identified variants. The sanitization procedure used here is inherently asymmetric, as bases are added or removed at only one end of the read. We make a distinction between a personal genome and the reference genome: the personal genome includes all of the SNPs and indels; the reference genome is the standard external metric. While the personal genome may not actually be constructed, it serves as a useful conceptual tool to understand the impact of the transformation involved in the sanitization procedure. We discuss three types of variants and the chosen sanitization method applied in the pBAM format:

##### SNP

A single-nucleotide variant/polymorphism is changed by mutating the variant to the reference allele in every read in which the mutation is observed.

##### Insertion

An insertion is sanitized by removing any fraction of the new inserted segment and adding the equivalent number of reference nucleotides to one end of the corresponding read.

##### Deletion

A deletion is sanitized by filling the reference nucleotides into the part of the deleted segment occurring on any read, and then removing the equivalent number of nucleotides from that read.

#### Definitions

The genome is indexed by discrete positions *i*. The coverage prior to and after sanitization are functions of the genomic position, and are labeled as *c^pre^* (*i*) and *c^post^* (*i*), respectively. The read length is fixed at *L_R_*. The size of the insertion is labeled 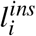, while the size of the deletion is labeled as 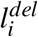, where in both cases the position *i* marks the start position of the indel. In addition, *N*(*i*) = the number of reads that start at position *i* in the mapping to the personal genome.

##### SNPs

Every read containing a SNP will be modified, with the alternate allele replaced by the corresponding reference allele. Under the assumption that the presence of this SNP does not alter the mapping efficiency (say, if the other variants within a particular read sum to *m^mis^*^−1^, then this SNP will lead to that read being dropped) and thus read dropout, we see that *c^pre^*(*i*) = *c^post^*(*i*). Therefore, no impact will be observed, unless one looks through the mapping quality control (QC) and finds all the reads overlapping a given locus have slightly lower quality. This might be possible, unless the QC metadata is being modified explicitly in the sanitization procedure (see Supplementary Information).

##### Short Indels

For indels, we consider the mapping changes due to the sanitization procedure in the following.

###### (1) Insertions

The variant is indexed by position *i*, where the insertion occupies the base pairs from *ℐ*+ 1 to 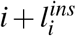. We consider the following cases:

- 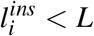: No individual read will dropout in this case due to the presence of the insertion. Consider a case in which the added nucleotides are on the end of a higher genomic position for the sake of clarity. The process of sanitization leads to an additional build-up of reads downstream of the insertion point. This happens due to the replacement process discussed above. Certain reads that would have been mapped to the insertion in the personal genome of the individual are now added downstream of the insertion in the mapping to the reference genome. This allows us to quantify the read build-up in terms of the start positions of the reads. Thus, for all reads that overlap with the insertion the following transformation occurs:

– 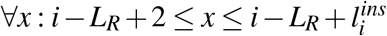, all reads starting at position *x* in the personal genome are newly mapped to the reference in the interval [*i*+ 1, *x* + *L_R_* + 1].
– 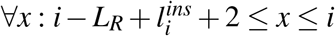, all reads starting at position *x* in the personal genome are newly mapped to the reference in the interval 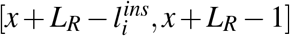.
– 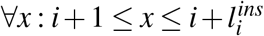, all reads starting at position *x* in the personal genome are newly mapped to the reference in the interval 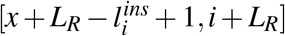. The genomic footprint of the sanitization procedure is the interval [*i*+ 1, *i* + *L_R_*]. The resultant read build-up of the sanitized BAM relative to the original BAM is thus given by the integral/discrete sum over all the accumulated contributions described above (again, *x* is the position along the personal genome, with the insertion in place; *α* in the equation below is the position along the reference genome in the downstream interval [*i*+ 1, *i* + *L_R_* − 1] that is impacted by read build-up due to post-sanitization remapping):

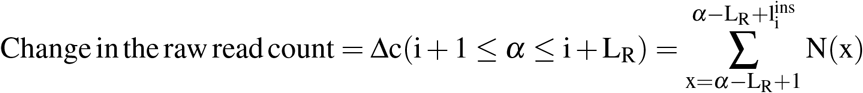 For example, imagine an ideal case where all the reads are uniformly distributed in the given region. This means that *c* (*x*) = constant = c and 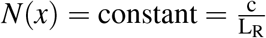 across the original mapping. Note that in the ideal case of uniformly distributed reads, the number of reads that begin at a locus is the total number of reads that overlap with a locus, divided by the length of each read. Thus,

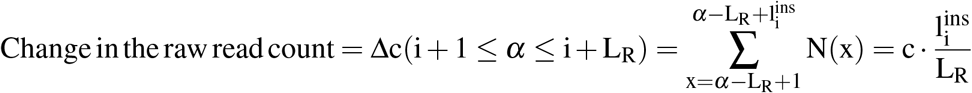 Thus, the fold-change in the coverage is given by 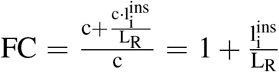 in the interval *i*+ 1 ≤ *α* ≤ *i*+ *L* If we define, FC(*α*) = exp(e_*α*_), we end up with 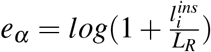 in the interval *i*+ 1 ≤ *α* ≤ *i*+ *L*. The general formula is

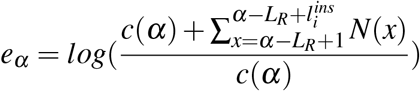
- 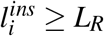: This case will be slightly different from the above case, as some of the reads will completely overlap with the insertion region and never contribute to the remapping build-up downstream. The calculation would then ignore the reads that overlap significantly with the insertion. We do not discuss this situation here, as these reads will have soft or hard clipping in their CIGARs (split reads) and will be treated differently as discussed above.

###### (2) Deletions

The variant is indexed by position *i*, where the deletion removes the reference base pairs from *i*+ 1 to 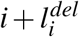. We exclusively consider 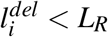 in this case, with the understanding that longer deletions would require slight modifications of the following calculations. The mapping to the reference genome after sanitization results in loss of coverage from regions downstream of the deletion, and an equal gain in regions of the deletion. The mapping changes are as follows:

- 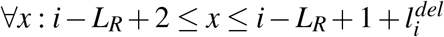, all reads starting at position *x* in the personal genome are removed from 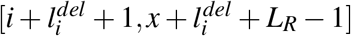 and newly mapped to the reference in the interval [*i*+ 1, *x* + *L_R_* − 1].
- 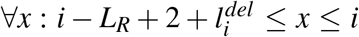, all reads starting at position *x* in the personal genome are removed from 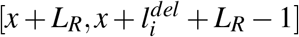 and newly mapped to the reference in the interval 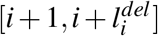.

The genomic footprint of the sanitization procedure is the interval 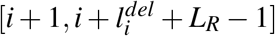. The change in coverage can be calculated in a manner similar to the case of insertions, with the additional notion that reads that are remapped to the deleted segment are drawn from downstream portions of the genome:

If the gain in the raw read count in the deleted segment of the reference genome is *Gain*, then

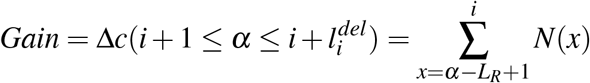

If the loss in the raw read count downstream of the deleted segment is *Loss*, then

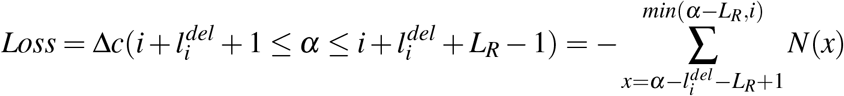

It is not possible to compute the fold change in the coverage in the deleted segment, as the coverage is 0 pre-sanitization. However, it can be calculated by adding a pseudo count to the coverage pre-sanitization. In the downstream segment, the fold change in the coverage is given by

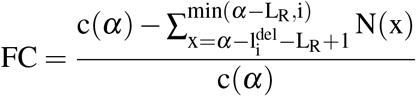

for 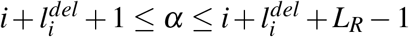 and with 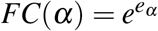, we have

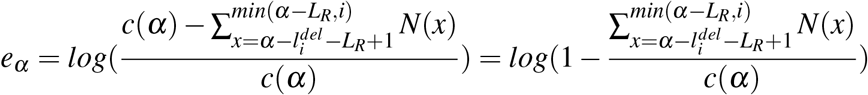

Again, considering the example of constant coverage discussed for the insertions, we have

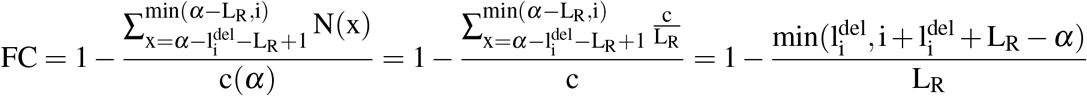

The upper bound on the fold change is given by 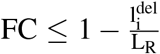. Note that the formula and the genomic footprint are different from those in the insertion case.

##### Suggestions for a higher utility pBAM while preserving privacy

After quantifying gene expression or other functional genomics quantities such as transcription factor binding enrichment from a BAM and corresponding pBAM file, one can pinpoint particular indels that overlap with the functional regions of the genome. We can then calculate the joint genotyping probability of these functional indels in a cohort of individuals (e.g., 1,000 Genomes individuals) in order to determine how rare or common they are. We can then systematically mask rare indels one by one until the difference between BAM and pBAM files are within noise levels (e.g., less than the difference between two replicates).

### 4.7 Functional genomics data processing

RNA-Seq data were processed using STAR [31] for alignment and RSEM [28] for quantification. ChIP-Seq data were processed using bwa [30] for alignment and MACS2 [34] for peak calling. ScRNA-Seq data were processed using HCA’s optimus pipeline [35] for quantifications and further analyzed using the Scanpy [36] (A jupyter notebook. The figures and the necessary input files can be found at privaseq3.gersteinlab.org).

## 5 Supplemental Information

**Figure S7:**
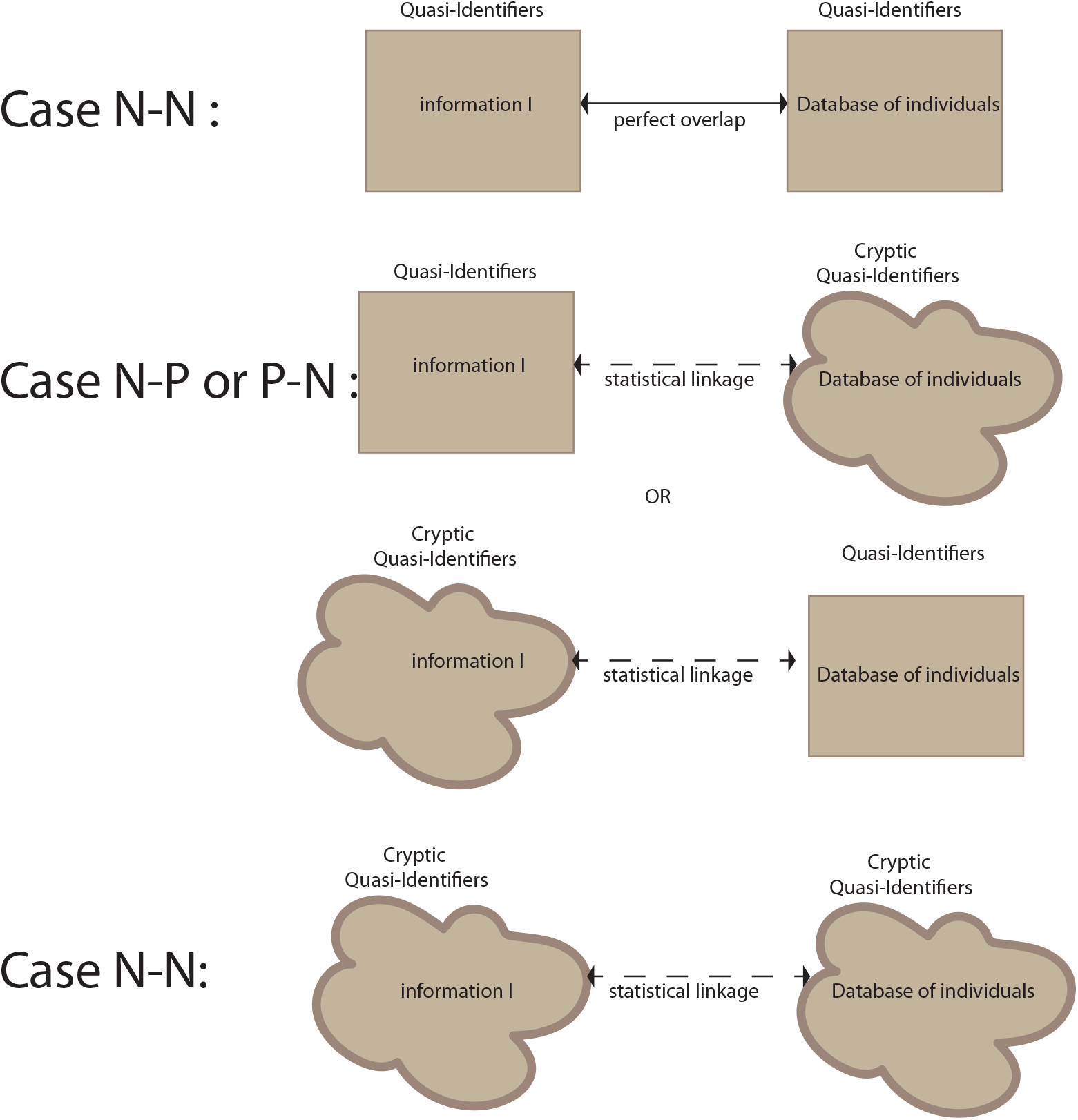
Different cases of linkage attacks.

**Table S1:** Mapping Statistics for WGS data.

| Sample Name | # of Total Reads | Percentage of Mapped Reads |
| --- | --- | --- |
| IndA-Cup1 | 127,850,309 | 82.36 |
| IndA-Cup2 | 137,723,180 | 82.14 |
| IndA-Cup3 | 160,138,265 | 61.50 |
| IndA-Cup4 | 215,379,662 | 75.98 |
| IndA-Cup5 | 157,769,539 | 83.56 |
| IndA-Cup6 | 121,329,654 | 83.54 |
| IndB-Cup1 | 207,414,742 | 83.58 |
| IndB-Cup2 | 151,139,298 | 62.65 |
| IndB-Cup3 | 122,164,133 | 83.76 |
| IndB-Cup4 | 220,846,681 | 77.93 |
| IndB-Cup5 | 140,335,759 | 85.26 |
| IndB-Cup6 | 172,391,527 | 88.97 |

**Table S1:** Genotyping statistics for different technologies.

|  | WGS |  |  | Genotyping Array |  |  |  | ONT |  |  |
| --- | --- | --- | --- | --- | --- | --- | --- | --- | --- | --- |
| samples | Called variants | Correct variants | Overlap with RNA-Seq | Called variants | Correct variants | Overlap with RNA-Seq | Call rate | Called variants | Correct variants | Overlap with RNA-Seq |
| indA-cup1 | 219,887 | 127,389 | 2,843 | 781,807 | 544,932 | 4 | 84% | 14 | 0 | 0 |
| indA-cup2 | 526,798 | 366,539 | 5,764 | 805,034 | 572,318 | 11 | 86% | 287 | 9 | 0 |
| indA-cup3 | 252,452 | 104,710 | 2,314 | 761,566 | 532,966 | 1 | 81% | 79 | 14 | 0 |
| indA-cup4 | 217,840 | 84,418 | 2,323 | 672,857 | 452,859 | 2 | 72% | 12 | 4 | 4 |
| indA-cup5 | 25,822 | 6,984 | 272 | 808,505 | 600,175 | 5 | 86% | 0 | 0 | 0 |
| indA-cup6 | 56,577 | 30,580 | 601 | 743,598 | 574,474 | 3 | 79% | 0 | 0 | 0 |
| indA-cup1 | 267,844 | 127,606 | 9,872 | 751,349 | 534,890 | 12 | 80% | 122 | 15 | 3 |
| indA-cup2 | 243,783 | 136,480 | 9,831 | 788,263 | 562,419 | 15 | 84% | 0 | 0 | 0 |
| indA-cup3 | 35,133 | 6,848 | 949 | 823,790 | 608,231 | 5 | 88% | 0 | 0 | 0 |
| indA-cup4 | 85,613 | 56,716 | 5,014 | 821,752 | 500,897 | 11 | 88% | 0 | 0 | 0 |
| indA-cup5 | 228,913 | 93,103 | 7,450 | 696,447 | 515,111 | 8 | 74% | 67 | 12 | 2 |
| indA-cup6 | 229,167 | 102,538 | 7,734 | 702,263 | 474,838 | 6 | 75% | 83 | 6 | 2 |

**Table S1:**
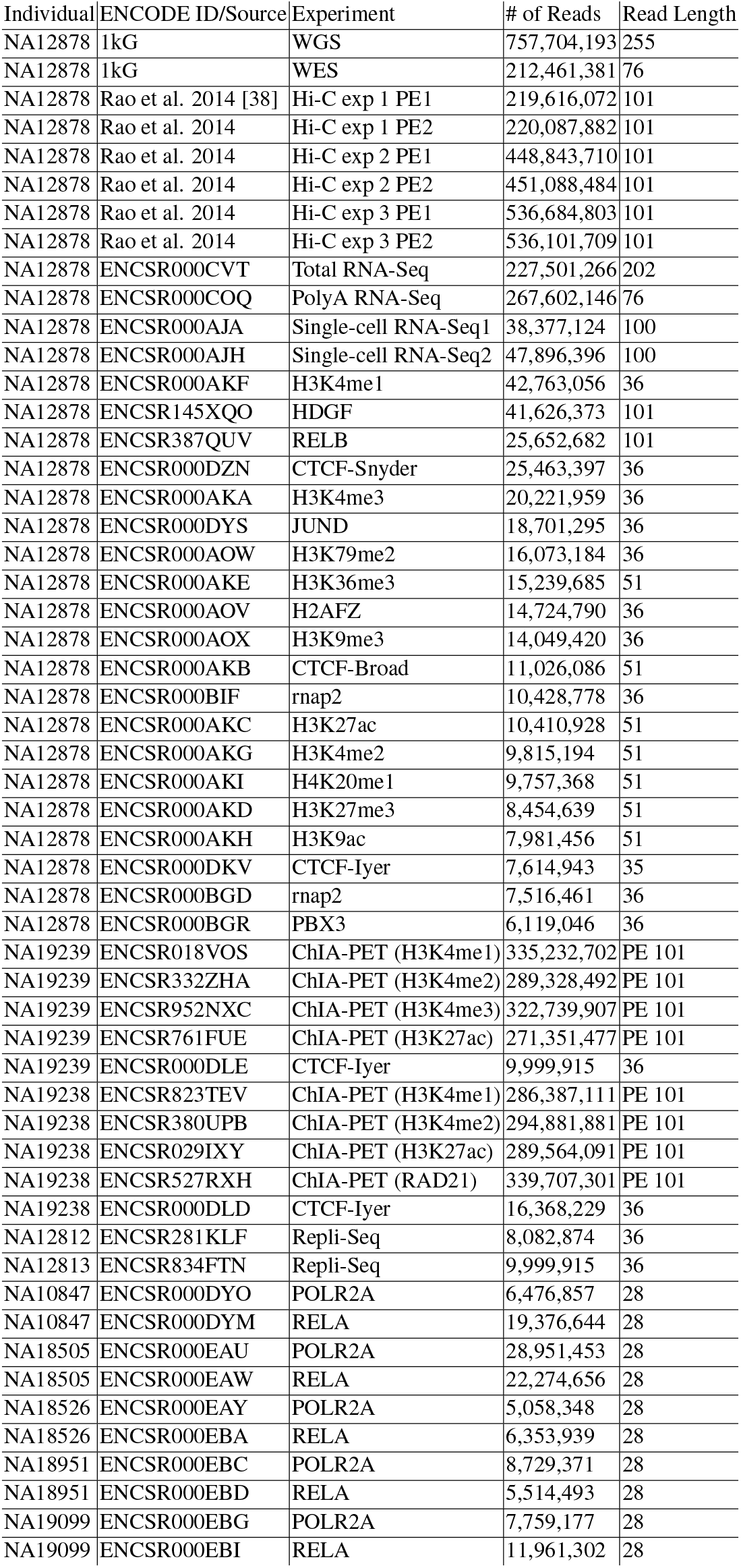
The functional genomics experiments used in this study with their total coverage

**Figure S7:**
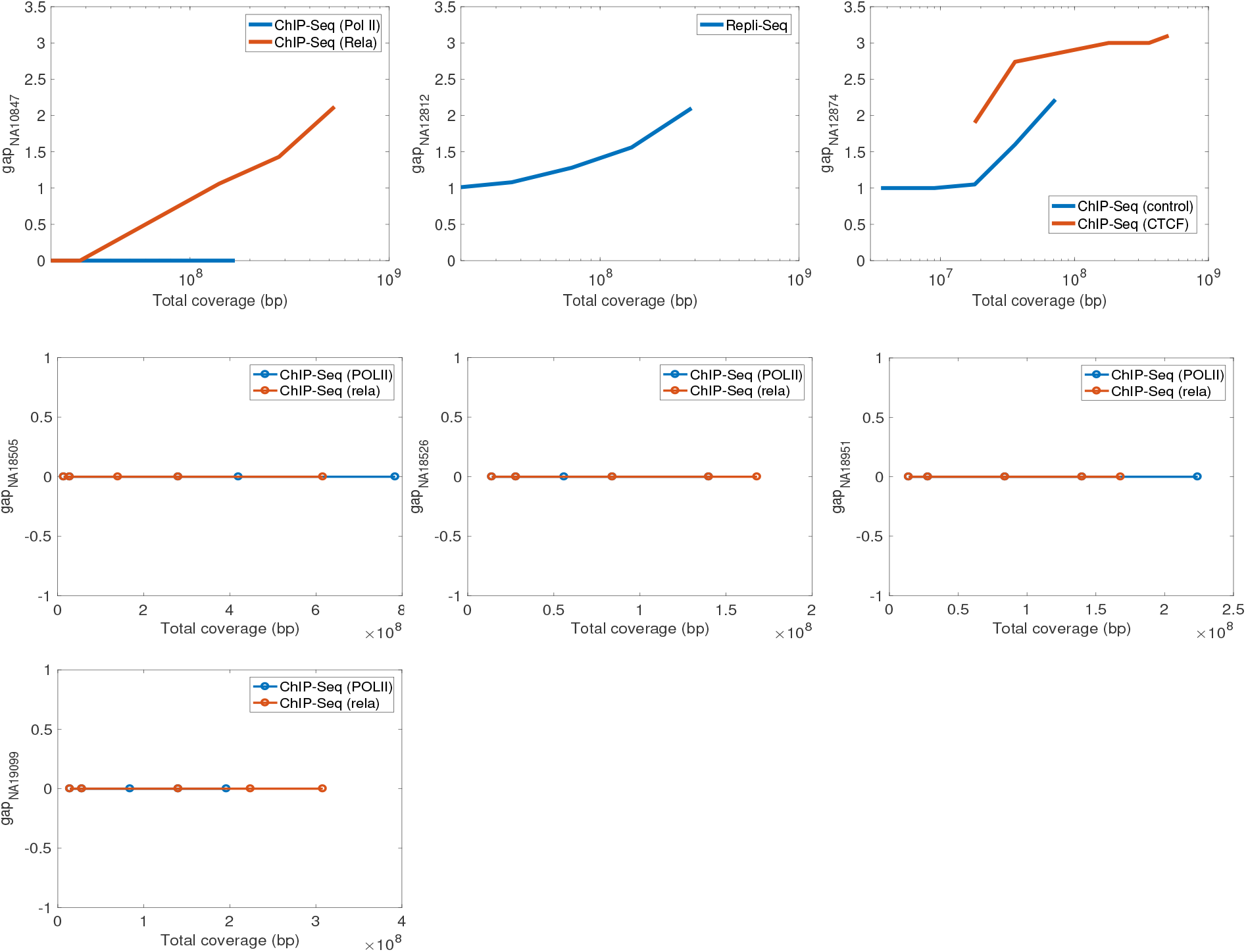
*gap* values for the remaining seven individuals with different functional genomics assays

**Figure S7:**
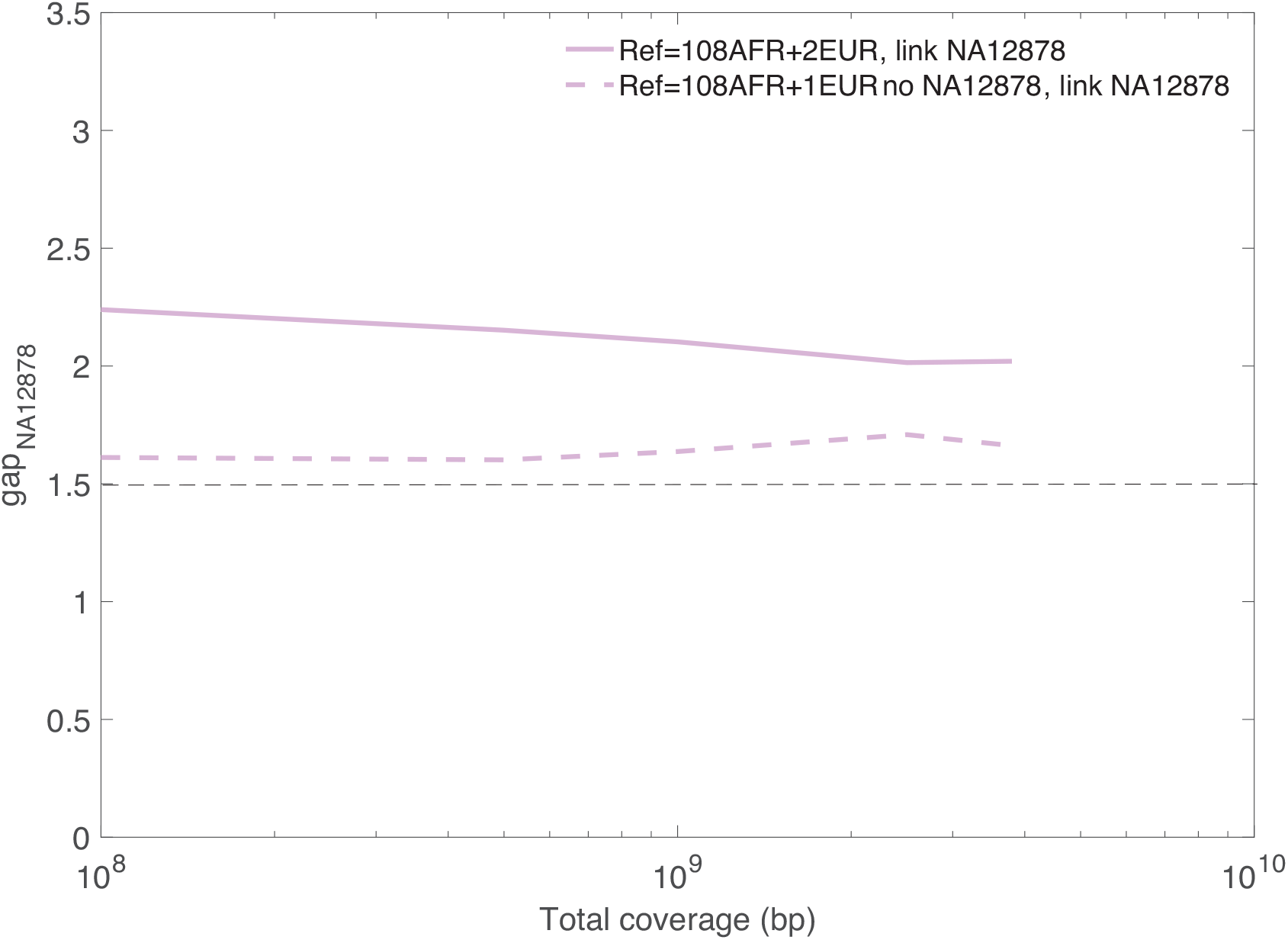
Linking NA12878 to a panel with 108 AFR individuals and one EUR individual, with and without NA12878 in the dataset.

### 5.1 Privacy-preserving file formats for functional genomics experiments

#### 5.1.1 Anonymizing the BAM files

We went through all the attributes of the BAM files and grouped them into two categories: (1) attributes to generalize with a common value and (2) attributes to remain as they are. The first category includes attributes that are leaking variants. They are the sequence, CIGAR attribute, and optional fields in the BAM files that are tagged with “AS” (alignment score), “MD” (string for mismatching positions), and “NM” (string for distance to reference). MAPQ values can also be revealing at times and are suggested to be sanitized in certain cases. The CIGAR gives information about how many matching and non-matching nucleotides there are in the read, with respect to a reference genome. As a result, one can call variants by looking at the non-matching nucleotides. We converted all the CIGARs to perfectly matching strings. For example, if the read length is 35 and the CIGAR is 14M1X15M, then the CIGAR is converted to 35M. AS reveals information about the number of matching positions in a read. An adversary can predict whether a read contains variants by looking at the alignment score and subtracting it from the read length. MD reveals information about the mismatching positions and deletions in the reads and their corresponding nucleotides. For example, if there is a nucleotide in the read that is “A” in the 15th position of a 30 bp read, and if the reference allele for this position is G, then the MD tag will be “MD:Z:14MA15M”, which directly reveals the variant position in the read. NM reveals how many bases are different than the reference, which in turn gives away how many SNPs there are in the read. We converted all the alignment scores to the read lengths and all the MD, AS, and NM tags to a perfectly matching string (for example “MD:Z:30M” for the example above). For the sequence attribute, we find the position of the read in the reference genome and replace the sequence attribute with the sequence in the reference. The rest of the attributes of the BAM files are designated as the second category and kept as they are.

#### 5.1.2 pBAM

Privacy-preserving file formats can be generated for SAM, BAM, and CRAM files. For simplicity, we will refer to the regular files as BAM and the privatized files as pBAM. The difference between the regular and the privatized files are in the fields of CIGAR, sequence, alignment score, the string for mismatching positions, and the string for the distance to the reference. Note that any optional field that leaks sensitive information about the sample can be manipulated. We focus on AS, MD, and NM tags throughout this paper, since they are the most obvious leakages, but a module to manipulate any other tag can easily be added to pTools.

Let us assume the read length for the sequencing experiment is 30, which is the total number of nucleotides in a fragment. Below is an itemized description of how CIGARs are converted to privatized CIGARs, along with examples:

##### CIGAR in non-intronic reads (i.e CIGARs with no ‘N”)

- The CIGAR for perfectly mapped reads is the read length followed by the letter “M”, indicating every nucleotide in the read is mapped to the reference human genome. This also means that there is no variant in this read (unless indicated in the MD tag). In this case, a regular BAM has “30M” in the CIGAR and pBAM has ‘30M” in the CIGAR as well.
- The CIGAR for reads that contain a mismatch is marked with the letter “X”. For example, if the 10th nucleotide in the fragment has a mismatch, then the CIGAR in the regular BAM becomes “9M1X20M”. This usually means that there is a SNP on the 10th nucleotide of the fragment. Since we know the start coordinate of the read from the regular BAM, an adversary can easily infer that there might be a SNP on the “*start* + 10”^*th*^ coordinate of the genome of the sample. To prevent that, we convert “9M1X20M” to “30M” in the pBAM file. This conversion does not add any noise to the calculation of depth since “*start* + 10”^*th*^ is sequenced, just as a different letter.
- The CIGAR for reads that contain a soft-clipping is marked with the letter “S”. For example, if the first five nucleotides are soft-clipped from the fragment, then the CIGAR becomes “5S25M”. The start coordinate reported is the beginning of the mapped nucleotides, which is the 6th nucleotide of the fragment. In this case, we report the CIGAR as “30M” and keep the start coordinate as it is. This is because soft-clippings can be due to a structural variant, insertion, or deletion. The associated noise with this conversion is that the coordinates between “*start* + 26” and “*start* + 30” gain extra depth.
- The above point also applies to reads with hard clippings, which are marked by the letter “H”. For example, if the 1st to 21st nucleotides are hard clipped from the fragment, then the CIGAR becomes “20H10M”. In this case, we report the CIGAR as “30M”, ignoring the hard-clipped nucleotides. The associated noise with this conversion is that the coordinates between “*start*” and “*start* + 20” gain extra depth.
- Note that some analysis pipelines such as ENCODE RNA-Seq and ChIP-Seq processing pipelines do not include these clipped reads in their analysis, in which case these reads can be filtered out entirely as they add the most noise to the signal after sanitization.
- The CIGAR for reads that contain an insertion is marked with the letter “I”. For example, if the 23rd to 30th nucleotide in the fragment is an insertion, then the CIGAR in the regular BAM becomes “22M8I”. Since we know the start coordinate of the read from the regular BAM, an adversary can easily infer that there is an insertion on the “*start* + 23”^*th*^ coordinate of the genome of the sample. To prevent that, we convert “22M8I” to “30M” in the pBAM file. The associated noise with this conversion is that the coordinates between “*start* + 22” and “*start* + 22 + 8” gain extra read, i.e., depth.
- The CIGAR for reads that contain a deletion is marked with the letter “D”. For example, if the 13th to 14th nucleotide in the fragment is a deletion, then the CIGAR in the regular BAM becomes “12M2D16M”. Since we know the start coordinate of the read from the regular BAM, an adversary can easily infer that there is an deletion on the “*start* + 12”^*th*^ coordinate of the genome of the sample. To prevent that, we convert “12M2D16M” to “30M” in the pBAM file. This conversion adds noise to the deleted coordinates as well as the last two nucleotides at the end of the read, as total read length has to be capped at 30. This also prevents signal profiles from leaking small deletions as the curve that corresponds to the deletion will look smooth based on its neighboring nucleotides.
- There are also CIGARs that may have multiple of the above letters. Here are a few examples and solutions:

##### Cigars in intronic reads (i.e., CIGARs with ‘N”)

- Cigars for perfectly mapped reads but split due to the introns are split by the letter “N”. For example, if there is a 1,000 nucleotide intronic region between mapped regions, it can have a CIGAR as “10M1000N20M”. In this case pBAM will have a CIGAR of “10M1000N20M” as well.
- If the reads are split in the mapped regions due to a mismatch, insertion, deletion, or clipping, then pBAM deals with them such that splice sites are as accurate as possible. Here are few examples;

– Cigar “3S15M1000N10M2D” becomes “18M1000N12M”, which does not add any noise to the splice site.
– Cigar “10M3D3M1000N3M2I9M” becomes “16M1000N14M”, which does not add any noise to the splice site.

Details of these examples are depicted in Figure S4.

**Figure S7:**
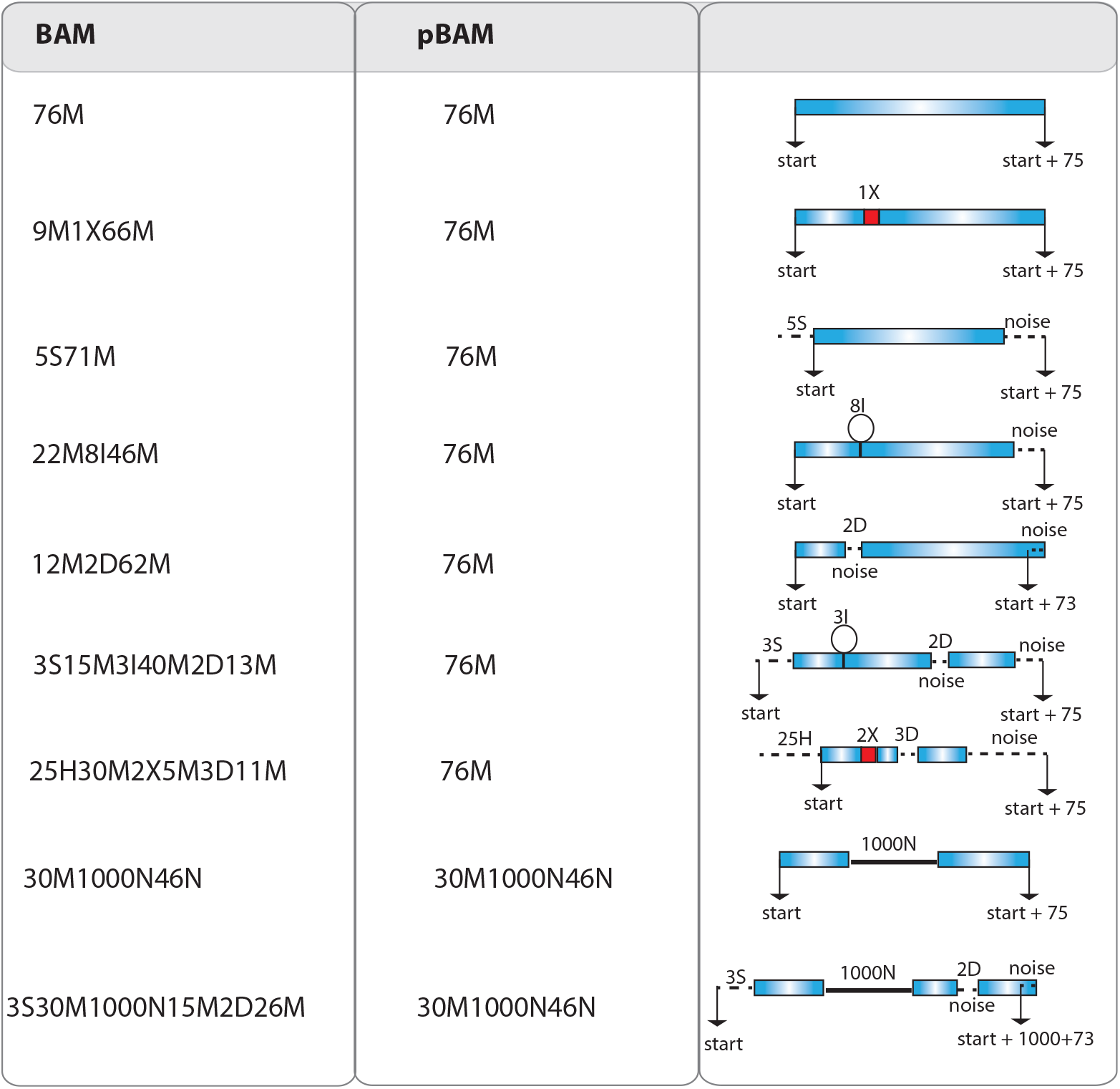
Visual representation of mapped fragments before and after converting the CIGARs to a pBAM file format. The insertions, deletions, soft and hard-clipping, and intronic reads are depicted. The noise that is added to the pBAM file to enhance privacy is also depicted in the fragments.

###### Transcriptome alignments

pTools searches the reference transcriptome for the position of the transcripts and reports the reference transcriptome sequences in the pBAM. We used the reference transcriptome files that are generated by STAR software [31].

#### 5.1.3 .diff files

.diff files contain the difference between the original BAM files and the pBAM files in a compact form. If the information is already available in the reference human genome such as sequence of the fragment, then the.diff file does not report it. This approach keeps the.diff files as small as possible. These files require special permission for access and contain the private information about the individual. To be able to go back and forth between BAM and pBAM files using the.diff files, the BAM and pBAM files must be coordinate sorted.

**Figure S7:**
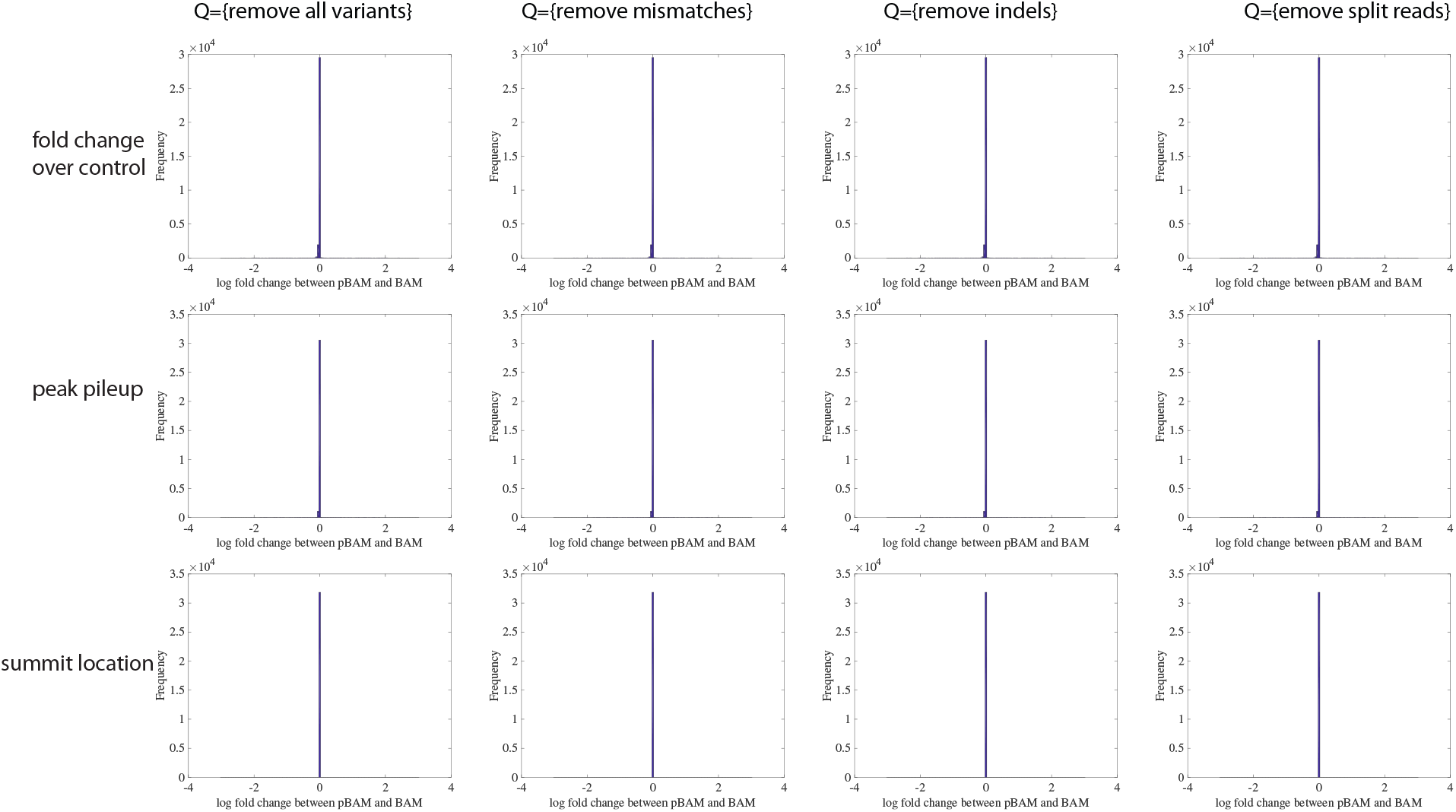
The difference between ChIP-Seq peak calling using BAM and pBAM as input for the fold change compared to control, the number of reads that pile up on the location of peak, and the location of the peak summit. Peak calling was performed using MACS2 [34].

### 5.2 Calculation of average and maximum leakage per variant

In the main text Discussion, we first overlapped the 1,000 Genomes variants with the exon annotations. We classified the variants into the following categories: exonic variants, exonic SNVs (excluding indels), exonic indels, and exonic small deletions. For each category, we calculated the self-information of the variant for all three possible genotypes (0, 1 and 2) as *h*(*s*_0_),*h*(*s*_1_) and *h*(*s*_2_). The average of self-information for each variant in each category is the average information leakage and the *max*(*h* (*s*^0^), *h*(*s*^1^), *h*(*s*^2^) is the maximum information leakage for that particular variant. We then calculated the mean and standard deviation for all the variants in each category. Total information leakage is calculated as the product of the total number of accessible variants and the average information leakage per variant. The distributions of the leakage can be seen in Figure S6 6. These distributions do not follow the normal distribution and are skewed, thus calculating the average is not the best approach. Therefore, mean leakage can be thought of an approximation that might be overestimated and since raw reads leak a lot more information than the signal profiles and eQTLs, the conclusion does not change with this approximation.

**Figure S7:**
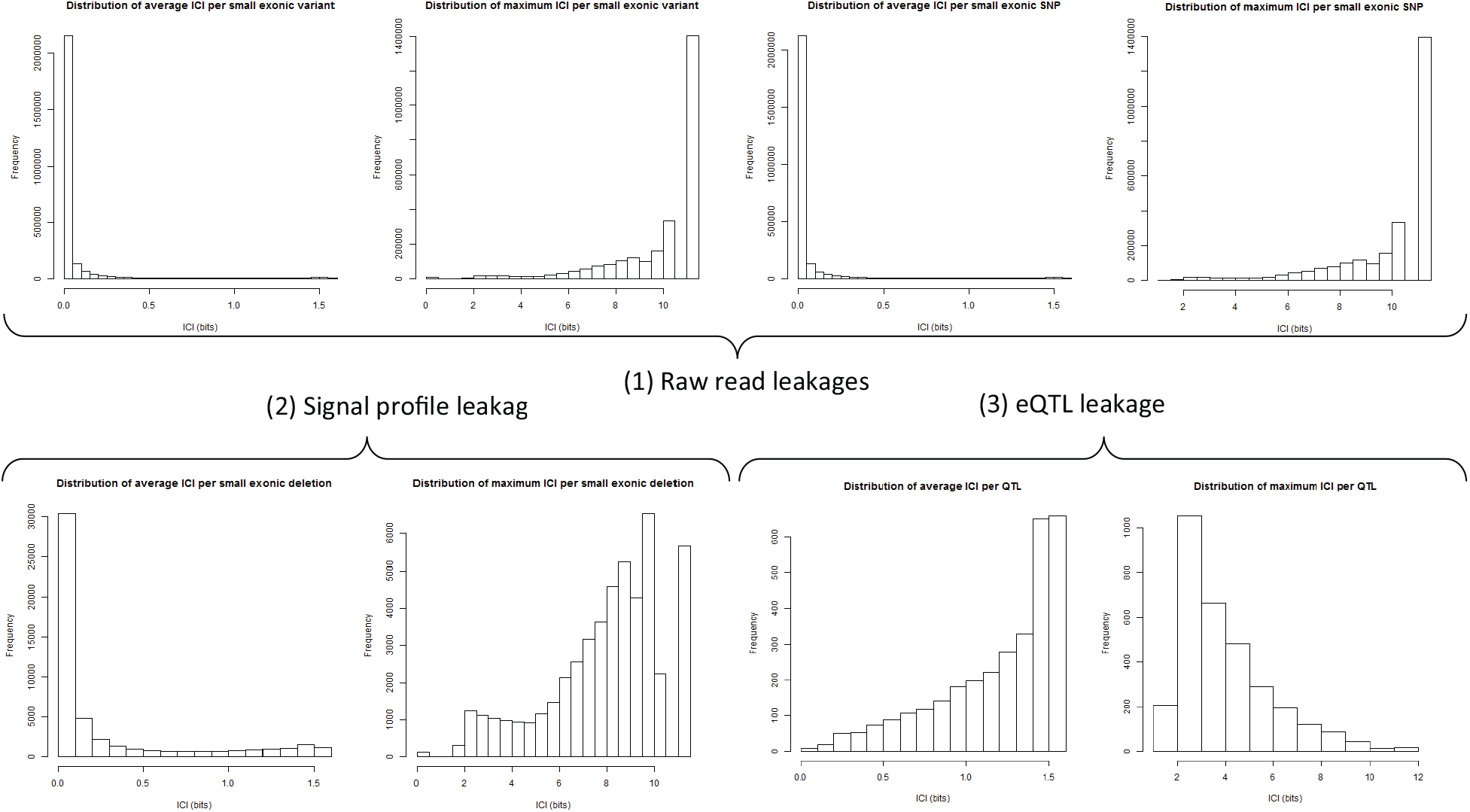
The distributions of the information leakage per variant in different levels of the data stack. Individual characterizing information (ICI) is calculated based on ref [13].

#### 5.2.1 Leakage from MAPQ

We found that reads with MAPQ values below the mean MAPQ contain insertions, deletions, and soft and hard clipping at a higher rate than expected (Figure 7), hence they might leak the location of large structural variants. In Figure S7, we analyzed a subsampled BAM file from a WGS dataset. The BAM files from functional genomics data are noisier than WGS, however the MAPQ values could still potentially be a source of variant leakage.

**Figure S7:**
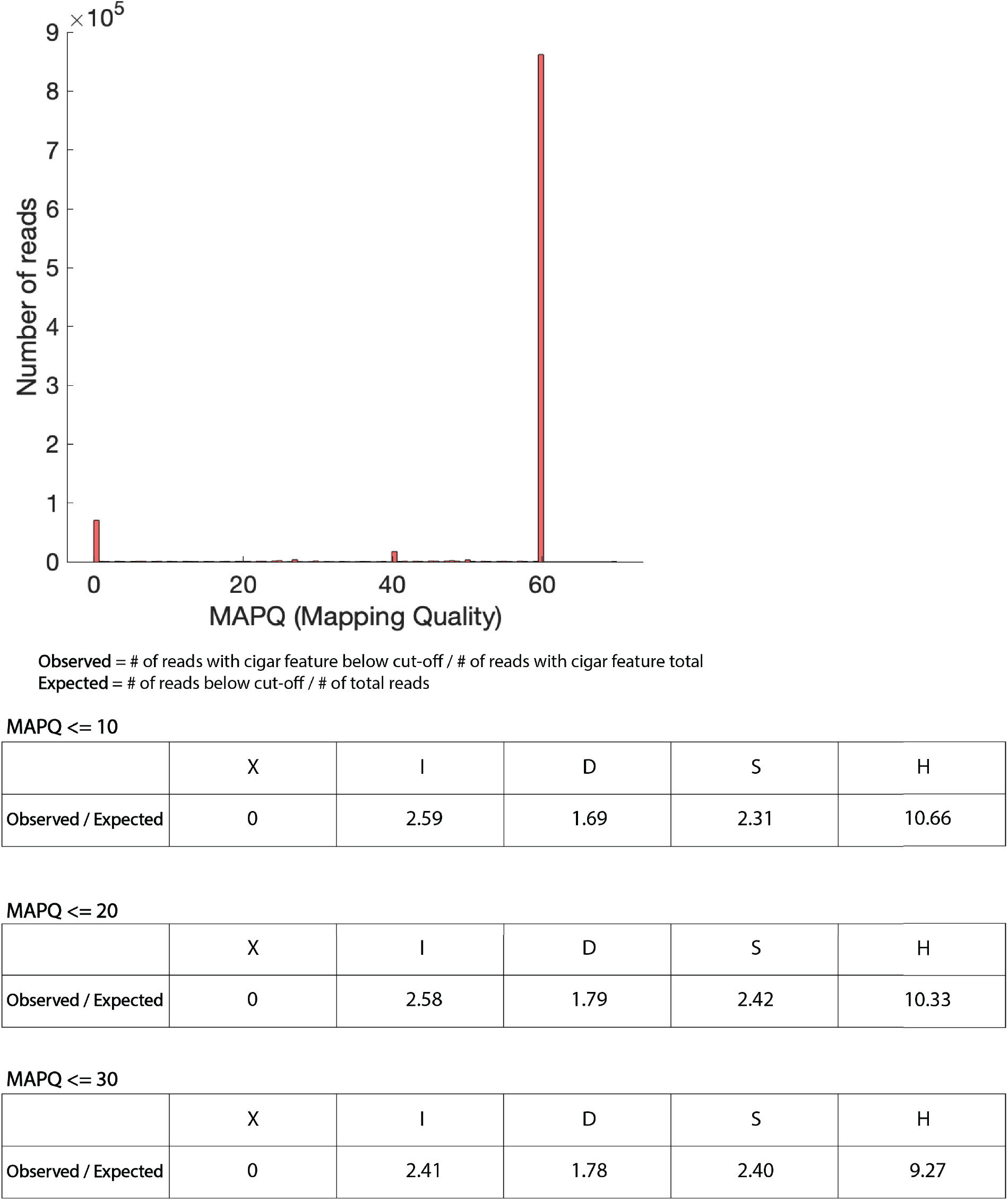
Potential variant leakage from MAPQ scores. As shown, the reads with potential large structural variants have smaller than expected MAPQ scores. An adversary can sort the MAPQ scores in a BAM file and guess the location of these structural variants that are mapped with low MAPQs.

